# The master energy homeostasis regulator PGC-1α couples transcriptional co-activation and mRNA nuclear export

**DOI:** 10.1101/2021.09.19.460961

**Authors:** Simeon R. Mihaylov, Lydia M. Castelli, Ya-Hui Lin, Aytac Gül, Nikita Soni, Christopher Hastings, Helen R. Flynn, Mark J. Dickman, Ambrosius P. Snijders, Oliver Bandmann, Heather Mortiboys, Sila K. Ultanir, Guillaume M. Hautbergue

## Abstract

PGC-1α plays a central role in maintaining the mitochondrial and energy metabolism homeostasis, linking external stimuli to the transcriptional co-activation of genes involved in adaptive and age-related pathways. The carboxyl-terminus encodes a serine/arginine-rich (RS) region and a putative RNA recognition motif, however potential RNA-processing role(s) have remained elusive for the past 20 years. Here, we show that the RS domain of human PGC-1α directly interacts with RNA and the nuclear RNA export factor NXF1. Inducible depletion of endogenous PGC-1α and expression of RNAi-resistant RS-deleted PGC-1α further demonstrate that the RNA-binding activity is required for nuclear export of co-activated transcripts and mitochondrial homeostasis. Moreover, a quantitative proteomics approach confirmed PGC-1α-dependent RNA transport and mitochondrial-related functions, identifying also novel mRNA nuclear export targets in age-related telomere maintenance. Discovering a novel function for a major cellular homeostasis regulator provides new directions to further elucidate the roles of PGC-1α in gene expression, metabolic disorders, ageing and neurodegenerative diseases.

## Introduction

The peroxisome proliferator-activated receptor gamma (PPARγ) coactivator 1 alpha (PGC-1α, also known as PPARGC1A) exerts a central regulatory role in sensing signal transduction pathways and maintaining the energy metabolism homeostasis in response to physiological processes during adaptive thermogenesis, exercise and fasting^1–4^. It is predominantly expressed in tissues with high energy demands such as heart, skeletal muscle, brown adipose tissue, liver, kidney and brain^1, 5^. Conversely, the expression level/ activity of PGC-1α and PGC-1α-dependent metabolic pathways are down-regulated in age and age-related diseases (muscle wasting, metabolic, neurodegenerative and neurodevelopmental disorders) ^6, 7^, highlighting the key importance of this master homeostasis regulator in human physiology.

PGC-1α co-activates the transcription of genes involved in mitochondrial biogenesis/maintenance and varied metabolic pathways through binding and activation of transcriptional activators, both nuclear receptors and other transcription factors, which co-activates the transcription of tissue-specific metabolic programmes^8^. PGC-1α was first discovered in brown adipose tissue as a mediator of non-shivering thermogenesis upon exposure to low temperatures^1^. There, it binds the nuclear receptors PPARα/ɣ^1, 9^ and retinoid X receptor-alpha (RXRα)^10, 11^ to co-activate the expression of uncoupling protein-1 (UCP-1) in mitochondria which uncouples the electron transport chain (ETC) to produce heat. In the liver and skeletal muscle interactions of PGC-1α with PPARs and RXRs lead to activation of the mitochondrial fatty acid oxidation while the co-activation of hepatocyte nuclear factor 4α (HNF4α) controls gluconeogenesis in the liver in an insulin-dependent manner^2, 3, 12^. Binding of PGC-1α to ERRα (estrogen-related receptor α) also regulates glucose and fatty acid metabolism in the striated muscle^13, 14^. On the other hand, PGC-1α is highly upregulated in skeletal muscles after cold exposure or exercise which leads to the increase of mitochondrial biogenesis and maintenance to meet energy demands^5^. In skeletal muscle and brown adipose tissue, PGC-1α directly induces the transcription of nuclear respiratory factors-1/2 (NRF-1/2). It further interacts and co-activates them to initiate in turn the transcription of nuclear-encoded mitochondrial transcription factor A (TFAM/mtTFA) ^5^ and other programmes of gene expression including electron transport chain and genes involved in oxidative phosphorylation (OXPHOS). Translocation of TFAM into the mitochondria subsequently stimulates replication and transcription of the mitochondrial genome promoting mitochondrial homeostasis and oxidative phosphorylation^15^.

Promoter recruitment of PGC-1α to nuclear receptors and transcriptional factors also stimulates interactions with chromatin remodelling factors^16^, and initiating/ elongating forms of the RNA polymerase II^17^ through binding with the general transcription initiation factor TFIIH^18^, the cyclin-dependent kinase complex CDK9^17^ or the Mediator^19^ to further promote the transcription of target genes. The expression and activity of PGC-1α is tightly regulated via methylation of its promoter^20, 21^, post-translational modifications^22–25^ and protein degradation^26–28^/ stabilisation^28, 29^.

In addition to the well-characterised transcriptional co-activator function of PGC-1α, the carboxyl-terminal domain harbours a Serine/Arginine (RS)-rich region flanked by a putative RNA Recognition Motif (RRM) reminiscent of SR-rich splicing factors, and a short motif which interacts with the CAP-binding protein 80 (CBP80) to promote the expression of a subset of transcripts^30^. PGC-1α is able to splice a reporter construct if recruited to a PGC-1α-regulated promoter^17^, suggesting it may co-transcriptionally splice transcripts it co-activates. However, it remains unknown whether PGC-1α plays a role in the splicing of cellular transcripts. On the other hand, the co-transcriptional recruitment of SR-rich splicing factors SRSF1,3,7^31–35^ and Aly/REF^36–39^ links deposition during pre-mRNA splicing and mRNA nuclear export through direct interactions with nuclear RNA export factor 1 (NXF1). This subsequently leads to the remodeling of NXF1 into a high RNA-affinity mode^40, 41^, which mediates transport through the channel of nuclear pores via transient interactions with protruding FG-repeats of nucleoporins^42, 43^. In 2020, PGC-1α was reported to differentially bind transcripts expressed from over a thousand genes in response to glucagon-induced fasting in mouse hepatocytes^44^. Although its role remains unknown, this study highlighted for the first time that the RNA-binding activity of PGC-1α is physiologically regulated.

Here, we show that the RS domain of PGC-1α directly interacts with poly-adenylated mRNAs and NXF1 in *in vitro* assays with purified recombinant proteins and RNA oligonucleotides as well as *in vivo* in human embryonic kidney (HEK) 293T cells. Using a stable inducible isogenic complementation system which allows the depletion of endogenous PGC-1α and the expression of an RNAi-resistant form of PGC-1α lacking the RS domain, we further demonstrate that the RNA-binding activity of PGC-1α is essential to the nuclear export of PGC-1α-activated mRNAs encoding proteins involved in the mitochondrial metabolism as well as to mitochondrial function and cell proliferation. In addition, PGC-1α is not involved in the binding or the nuclear export of non-transcriptionally co-activated target transcripts, further indicating that its nuclear mRNA export function is coupled to transcriptional co-activation, at least for the transcripts investigated in this study. A quantitative proteome analysis of the RNA/NXF1-binding mutant confirmed alterations of the RNA transport and mitochondrial respiratory oxidative metabolisms. This investigation also identified novel mRNA nuclear export targets involved in DNA repair and telomere maintenance, pathways directly relevant to the age-related functions of PGC-1α, but yet unknown to be regulated in a PGC-1α-dependent manner. Taken together, we discovered that PGC-1α, a major physiological regulator extensively studied over the past 20 years (>4,500 publications in *PubMed*), exhibits a novel mRNA nuclear export activity, essential to its function, at the heart of energy metabolism homeostasis.

## Results

### PGC-1α binds RNA via the RS domain

The carboxyl-terminal domain of PGC-1α harbours a serine-arginine-rich (RS) region flanked by a putative RNA recognition motif (RRM) and a CBP80-binding motif (CBM). To assess the potential RNA-binding ability of PGC-1α, we performed ultra-violet (UV) irradiation mediated covalent crosslinking of interacting RNA probes with purified recombinant proteins. PGC-1α full length could not be expressed in *E. coli* despite fusion to various solubility and expression tags including protein G domain B1 (GB1) and glutathione-S-transferase (GST). We further engineered a series of plasmids expressing GB1-hexa-histidine-tagged (GB1-6His) transcriptional co-activation (D1, aminoacids aa1-234), suppression/regulatory (D2, aa254-564), putative RNA-binding (D3, aa565-798), RS (aa565-633) and RRM+CBM (aa-634-798) domains of human PGC-1α (**Fig. 1a**). Recombinant proteins were successfully expressed in *E. coli* prior to ion metal affinity chromatography purification from soluble and insoluble fractions (**Supplementary Fig. 1**). Most domains, except D1 and CBM, were produced in inclusion bodies and were therefore solubilised using urea prior to large batch affinity purification in buffers with high salt concentrations to further disrupt potential contaminant interactions with bacterial RNA and proteins. Rapid dilution of denatured protein domains in the final binding assay buffers further allowed bypassing the intermediate concentrations of denaturing urea that typically lead to formation of intermolecular aggregation and misfolded proteins^45, 46^ (**Methods**). Purified human recombinant GB1-6His-tagged PGC-1α D3 or Aly/REF (a known general mRNA nuclear export adaptor^36, 47^) and 6His-MAGOH (a control protein which does not bind RNA^48, 49^) were then incubated in the same buffer conditions with ^32^P-radiolabelled synthetic AU-rich (AAAAUUx5) or GC-rich (GGGGCCx5) RNA probes prior to irradiation with UV where indicated (+). PhosphoImages showed that covalently bound RNA molecules remained associated with Aly/REF and PGC-1α D3 which can be visualized on the Coomassie-stained panels during the SDS-PAGE electrophoresis. The interactions were specific since no binding of RNA was detected in absence of UV-irradiation or with the negative control protein MAGOH (**Fig. 1b**). A similar investigation of D3 subdomains demonstrated that the RS domain is required and sufficient for direct binding to RNA (**Fig. 1c**). It is noteworthy that the addition of urea during the purification is not expected to affect potential folding of the RS domain as these arginine-serine-rich regions are intrinsically-disordered^50^ (**Supplementary Fig. 2**).

**Figure 1.**
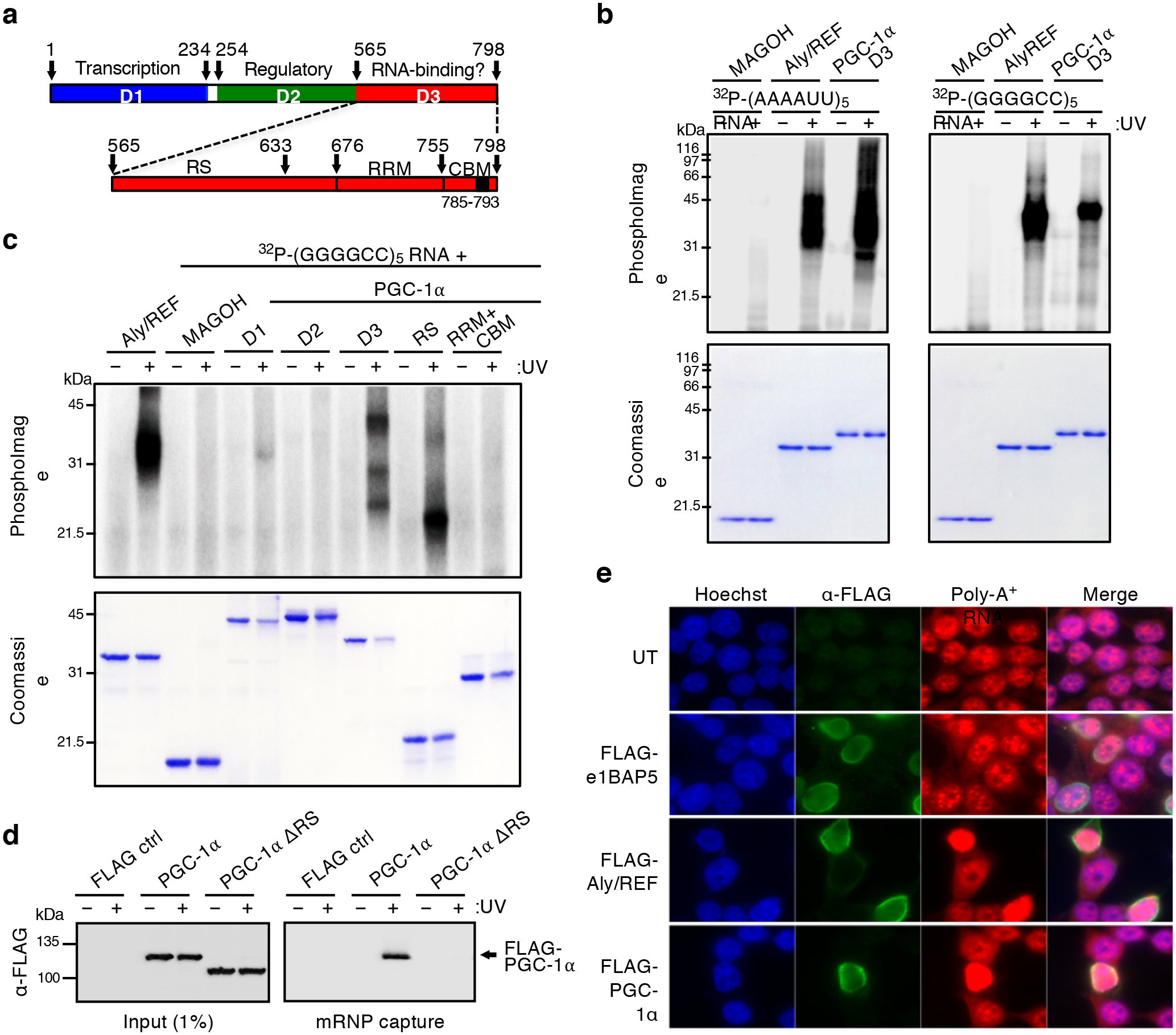
The RS domain of PGC-1α interacts with transcriptionally co-activated mRNAs. (**a**) Schematic representation of the primary structure of PGC-1α including the N-terminal transcriptional co-activation domain (blue), suppression/regulatory domain (green) and the putative C-terminal RNA-binding domain (red), which is further expanded below to highlight the RS, RRM and CBM domains. **(b)** Protein:RNA UV-crosslinking assays. Recombinant PGC-1α D3 (aa565-798) was incubated with AU- or GC-rich ^32^P-labelled RNA probes prior to UV irradiation (+) or not (-). Coomassie-stained gel shows recombinant proteins and phosphoimage shows radiolabelled RNA. Aly/REF and Magoh are included as positive and negative controls, respectively. **(c)** Protein:RNA UV-Recombinant PGC-1α domains were incubated with GC-rich ^32^P-labelled RNA probe prior to UV irradiation (+) or not (-). Coomassie-stained gel shows recombinant proteins and phosphoimage shows RNA. Aly/REF and Magoh are included as positive and negative controls respectively. **(d)** mRNP capture assay. Human HEK293T cells were transfected for 48 h with either FLAG control, FLAG-tagged PGC-1α wildtype or FLAG-tagged PGC-1α ΔRS. Cells were subjected to UV irradiation (+) or not (-). Protein extracts were subjected to denaturing mRNP capture assays using oligo-dT beads. Eluted proteins were analysed by western blotting probed with α-FLAG antibody. **(e)** Bulk poly(A)+ cellular distribution upon overexpression of PGC-1α. HEK293T cells were transfected with E1B-AP5/hnRNPUL1 (negative control), Aly/REF (positive mRNA export adaptor control) and PGC-1α for 48 hours followed by a 2 h treatment with 5 μg/ml Actinomycin D. Cells were stained with anti-FLAG antibody (green) to visualise transfected cells and Cy3-oligo(dT) (red) that anneal to the poly(A) tails of RNA molecules.

To evaluate the *in vivo* relevance of these findings, we performed UV-mediated protein:RNA crosslinks on live cells prior to denaturing messenger-ribonucleoprotein (mRNP) capture assays using oligo-dT beads. These showed that FLAG-tagged PGC-1α, but not the full-length protein lacking the RS domain (ΔRS), directly interacts with the fraction of poly(A)^+^-enriched mRNAs in human embryonic kidney (HEK293T) cells (**Fig. 1d**, **Supplementary Fig. 3**). We have previously shown that overexpression of the conserved mRNA nuclear export adaptor Aly/REF blocks the bulk NXF1-dependent nuclear export of poly(A)^+^ RNAs^51, 52^, likely due to excessive adaptor:NXF1 interactions which stall the high RNA-affinity remodeling of NXF1 in absence of recruitment to spliced transcripts. Similarly, Cy3-oligo-dT Fluorescence *In Situ* Hybridisation (FISH) experiments showed that FLAG-tagged Aly/REF or PGC-1α triggers nuclear accumulation of poly(A)+ RNA in anti-FLAG-stained cells highly overexpressing these proteins, while the control NXF1-binding ribonucleoprotein hnRNPUL1/E1B-AP5^52^ did not (**Fig. 1e**).

Taken together, our data demonstrate that the RS domain of PGC-1α directly interacts with RNA oligonucleotides as well as poly-adenylated RNA in human cells in agreement with a recent genome-wide iCLIP study reporting binding of PGC-1α to RNA in absence of strict consensus elements^44^. Similarly, we have previously reported that arginine-rich regions adjacent to RRM domains constitute broad-specificity RNA-binding sites for the nuclear mRNA export adaptors SRSF1,3,7 and Aly/REF^33, 34, 40, 53, 54^.

### The RS domain of PGC-1α interacts with NXF1

Our structural and functional investigations of the nuclear mRNA export adaptors SRSF1,3,7 and Aly/REF previously showed that NXF1 binds arginine residues flanked by prolines in short unstructured peptides^33, 34, 40, 53, 54^. Interestingly, short peptide sequences in the RNA-binding domain of PGC-1α exhibit similar characteristics (**Fig. 2a**). To assess potential interactions between PGC-1α and NXF1, we performed pull-down assays using bacterially-expressed GST-NXF1 and ^35^S-radiolabelled PGC-1α proteins synthesised in rabbit reticulocytes in the presence of RNase to obliterate indirect interactions bridged by RNA molecules^33, 34, 40, 51^. GST-NXF1 was co-expressed with p15/NXT1, a co-factor which improves its stability through heterodimerization with the NTF2-like domain (aa371-551) of NXF1 ^55, 56^. GST pull-down assays showed that PGC-1α full length (WT) and carboxyl-terminal D3 encompassing the RS region interact with NXF1 in an RNA-independent manner (**Fig. 2b**). It is challenging to detect the endogenous interaction of NXF1 with nuclear export adaptors in mammalian cells due to the transient nature of the NXF1-driven nuclear export process. Therefore, as in our previous reports, we co-transfected HEK293T cells with FLAG-PGC-1α wild type (WT) and Myc-NXF1 plasmids, showing specific interactions in RNAse-treated co-immunoprecipitation assays (**Fig. 2c**). It is noteworthy that FLAG-tagged PGC-1α constructs were reported to have moderate overexpression in other studies^17, 26^. The transcription-export (TREX) complex, which is deposited during splicing^37–39^, provides a platform for the high RNA-affinity remodelling of NXF1 and its recruitment along spliced mRNA molecules^39–41^. Co-immunoprecipitation assays in HEK cells showed that several endogenously-expressed subunits of TREX (THOC1, UAP56, Aly/REF) co-immunoprecipitates with PGC-1α in an RNAse-independent manner (**Fig. 2c**). Interestingly, the association of PGC-1α and TREX is stabilised in cells transfected with the RNA-binding mutant (ΔRS), suggesting a stalling of the dynamic mRNA nuclear export process which requires handover of the processing mRNA from the nuclear export adaptor to NXF1^40, 41^ (**Fig. 2c**). In agreement with the biochemical data, we also observed that endogenous NXF1 and PGC-1α proteins partially co-localise in the nuclei of HEK293T cells (**Fig. 2d**, **Supplementary Fig. 4**).

**Figure 2.**
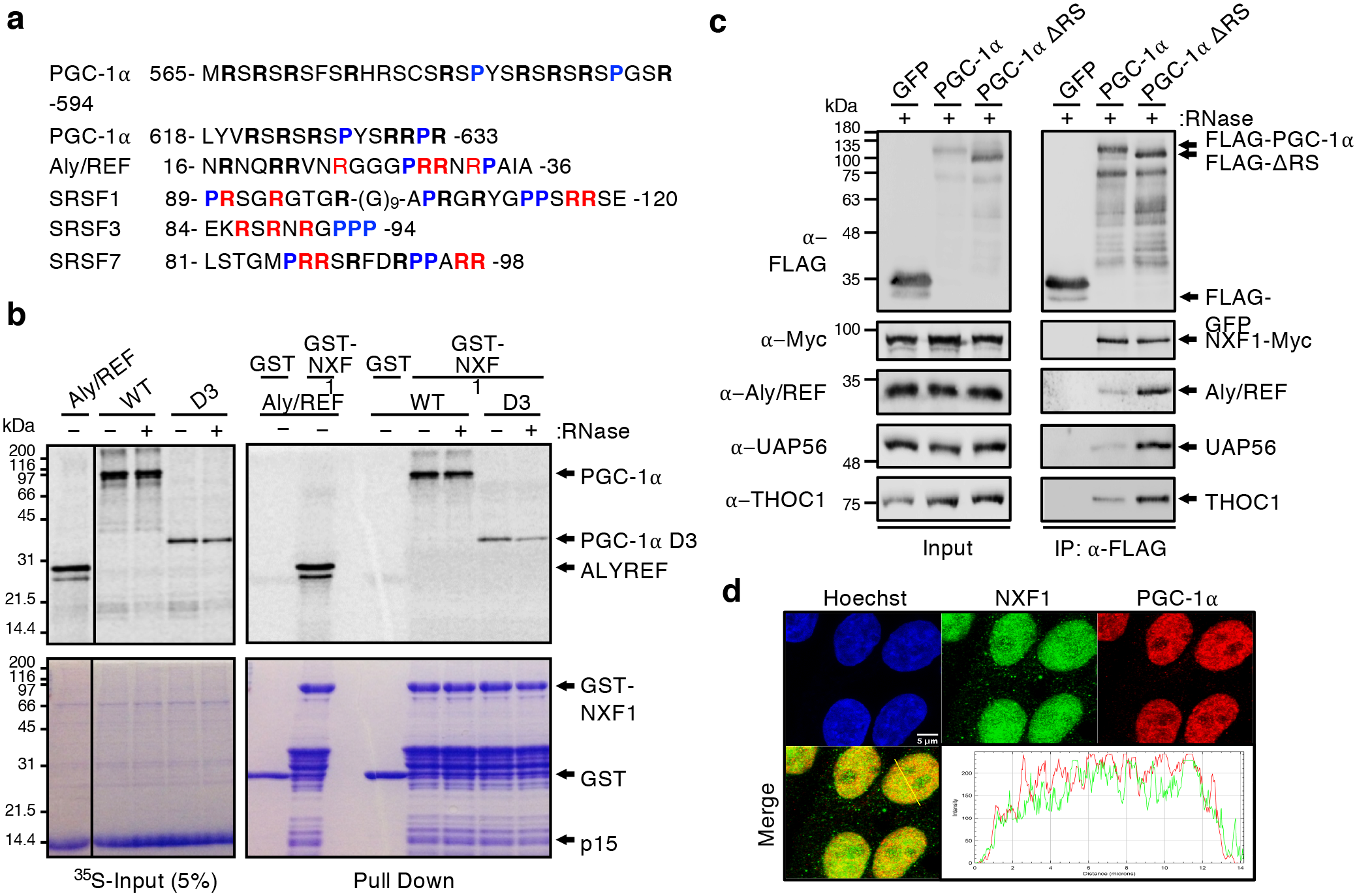
PGC-1α binds the nuclear export receptor NXF1 and associates with the TREX complex. (**a**) Schematic showing arginine residues flanked by prolines in short unstructured peptide sequences of the arginine-rich RNA-binding domains of PGC-1α, Aly/REF, SRSF1, SRSF3 and SRSF7. Arginine residues labelled in red were previously shown to interact with RNA. **(b)** GST pull down assay. PGC-1α full length (WT) and D3 were expressed and radiolabelled with ^35^S-methionine in rabbit reticulocyte lysate prior to affinity purification with GST or GST-NXF1:p15 bound to glutathione-coated beads. Aly/REF was used as a known mRNA export adaptor control. Coomassie-stained gels (bottom) show input and pull-down with immobilised GST and GST-NXF1:p15. PhosphoImages (top) show expression of Aly/REF and PGC-1α (input) and co-purified proteins (Pull Down). The interaction of PGC-1α with GST-NXF1:p15 is not ablated by RNase treatment. **(c)** Co-immunoprecipitation assays. HEK293T cells were co-transfected with plasmids expressing FLAG-tagged PGC-1α WT, ΔRS or GFP as a control and c-Myc-tagged NXF1. Whole-cell lysates were subjected to anti-FLAG immunoprecipitation. Western blot was carried out with antibodies for FLAG, c-Myc, Aly/REF, UAP56 and THOC1. Left panel represents input lanes. Right panel shows FLAG IP and co-immunoprecipitation of TREX proteins. **(d)** Confocal immunofluorescence microscopy in HEK293T cells. NXF1 and PGC-1α antibodies were used to label endogenous NXF1 and PGC-1α proteins in green and red, respectively. Bar scale: 5 µm. Green and red pixel intensities were quantified alongs a nuclear cross-section highlighted with a yellow line.

We have previously shown that the hallmark of mRNA nuclear export adaptors is the presence of short unstructured arginine-rich regions with overlapping RNA/NXF1-binding sites^33, 34, 40, 53, 54^. Since the RS domain of PGC-1α directly bind RNA (Fig. 1), we sought to test whether it would also interact with NXF1 (**Fig. 3a**). RNAse-treated GST pull down assays using ^35^S-radiolabelled PGC-1α protein domains synthesised in rabbit reticulocytes showed that ALY/REF, PGC-1α D3 and RS domains specifically interact with NXF1 (**Fig. 3b**). Similarly, we show that the RS domain is required and sufficient for the direct binding of NXF1 using recombinant stringently-purified protein domains expressed in *E. coli* (**Fig. 3c, Methods**). Interestingly, D3 interacts with NXF1 when synthesised in a mammalian transcription/translation-coupled system which allow for post-translational modifications (PTMs) (Fig. 3b) whereas it does not when expressed in bacteria which do not support mammalian PTMs (Fig. 3c), suggesting that PTMs play a role in the NXF1-dependent regulation binding of PGC-1α. However, preliminary experiments using phosphatase or methyl transferase inhibitors did not allow elucidating this further. As expected, deletion of the RS domain in the full length PGC-1α expressed in rabbit reticulocytes considerably attenuated the interaction with NXF1 (**Fig. 3d**). Taken together, we show that the RS domain of PGC-1α is required for interactions with NXF1 and RNA both *in vitro* and *in vivo* in human cells.

**Figure 3.**
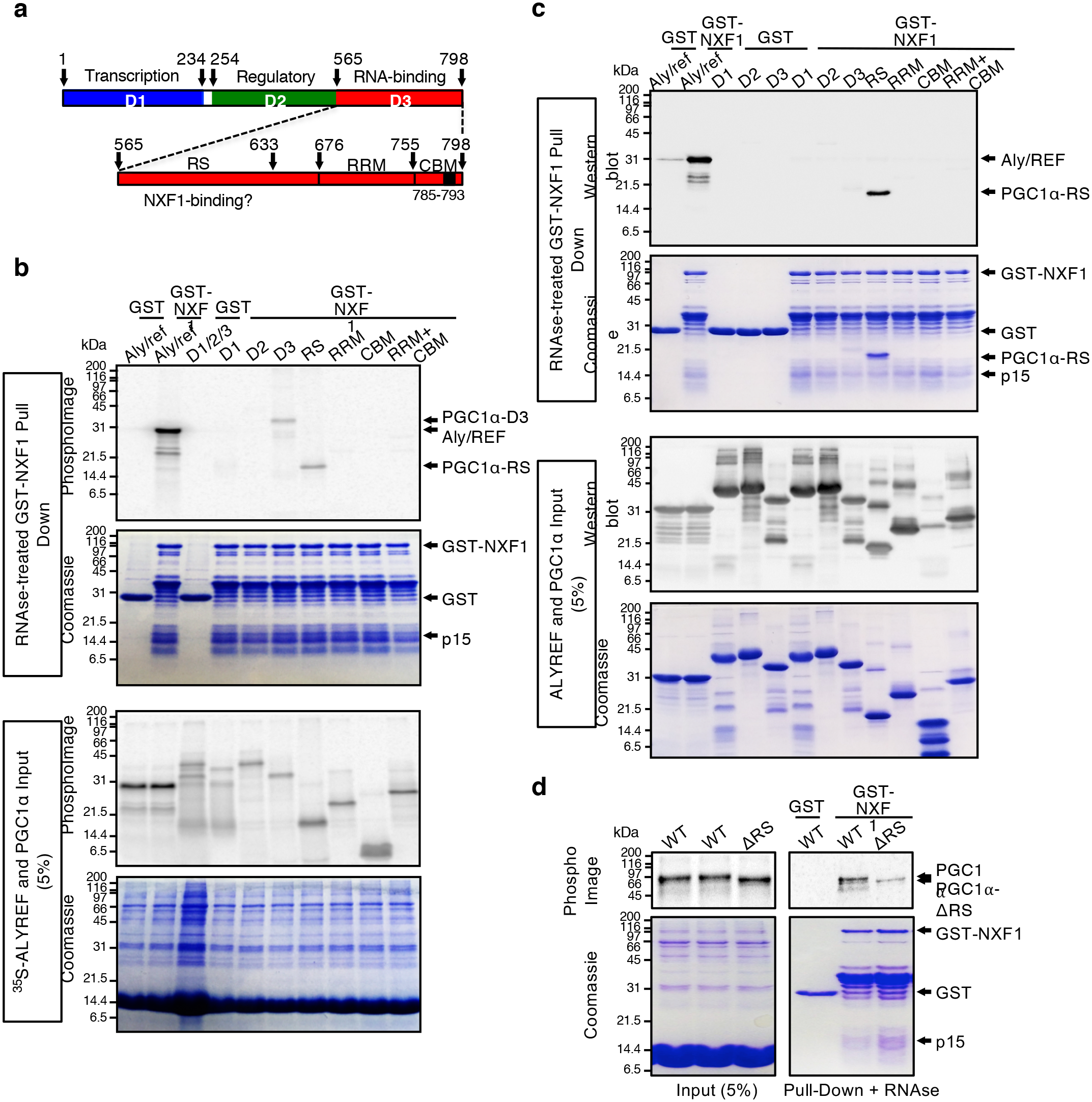
The RS domain of PGC-1α binds NXF1. (**a**) Schematic representation of the structure of PGC-1α including the N-terminal transcriptional activation domain (blue), the middle suppression domain (green) and the C-terminal domain (red) which is further expanded below to highlight the RS, RRM and CBM domains. Putative NXF1 binding site is highlighted. **(b)** GST pull down assays. PGC-1α protein domains were expressed and radiolabelled with ^35^S-methionine in rabbit reticulocyte lysate prior to affinity purification with bacterially-expressed GST or GST-NXF1:p15 bound to glutathione-coated beads. Aly/REF was used as a known mRNA export adaptor control. Bottom 2 Coomassie-stained gel and PhosphoImage show input lanes with ^35^S-radiolabelled Aly/REF and PGC-1α protein domains expressed in rabbit reticulocytes. Top 2 Coomassie-stained gel and PhosphoImage show immobilised GST and GST-NXF1:p15 with co-purified proteins. Aly/REF, PGC-1α D3 and PGC-1α RS bind NXF1. All binding reactions were treated with RNase. **(c)** GST pull down assays using bacterially-expressed and stringently-purified recombinant proteins. PGC-1α protein domains were expressed in *E. coli* prior to affinity purification with bacterially-expressed GST or GST-NXF1:p15 bound to glutathione-coated beads. Bottom 2 Coomassie-stained gel and α-6His western blot show input lanes with ALY/REF and PGC-1α protein domains. Top 2 Coomassie-stained gel and α-6His western blot show immobilised GST and GST-NXF1:p15 with co-purified proteins. All samples were treated with RNase. **(d)** GST pull down assays. Bacterially-expressed GST or GST-NXF1:p15 bound to glutathione-coated beads were incubated with ^35^S-PGC-1α WT and ΔRS radiolabelled in rabbit reticulocytes in presence of RNAse. Coomassie images show input and pull-down with immobilised GST and GST-NXF1:p15. PhoshoImages show ^35^S-radiolabelled PGC-1α input (left) and pulldown (right) lanes.

### Investigating the physiological relevance of the potential mRNA nuclear export activity of PGC-1α

To assess the functional importance of the RS domain, we sought to engineer human cell complementation systems which allow the depletion of endogenous PGC-1α and its replacement with either a control FLAG-tagged PGC-1α wildtype protein or the mutant lacking the RS domain. For this purpose, stable isogenic HEK293T-FlpIn cell lines were generated. They allow doxycycline-inducible expression of the following transgenes integrated at the single genomic FRT site (Flp Recognition Target): (i) Control-miRNA (Ctrl-RNAi) which does not target human transcripts; (ii) PGC-1α miRNA cells (PGC1α-RNAi) expressing 2 chained miRNAs recognising nucleotide sequences in D1 and D2 of PGC-1α; (iii) RNAi-resistant FLAG-tagged full-length PGC-1α cells co-expressing the PGC1α-RNAi cassette (WT-res); (iv) RNAi-resistant FLAG-tagged RS-deleted PGC-1α cells co-expressing the PGC1α-RNAi cassette (ΔRS-res). Control cells with recombination of an empty plasmid at the FRT site also serve as a control healthy HEK293T cell line (Sham) (**Fig. 4a**). In contrast to Ctrl-RNAi cells, doxycycline-induced expression of the PGC1α-RNAi cassette leads to robust and specific depletion of endogenous PGC-1α transcripts and protein respectively quantified by qRT-PCR (**Fig 4b**) and western blot using an anti-PGC-1α antibody validated in previous studies^44, 57, 58^ (**Fig. 4c-d**). Both the depletion of endogenous PGC-1α and expression of FLAG-tagged PGC-1α wildtype protein or mutant lacking the RS domain were confirmed in the WT-res and ΔRS-res cell lines (**Fig. 4e-f**). These data indicate that these cell lines allow inducible depletion of PGC-1α and/or co-expression of RNAi-resistant PGC-1α and PGC1α-ΔRS proteins.

**Figure 4.**
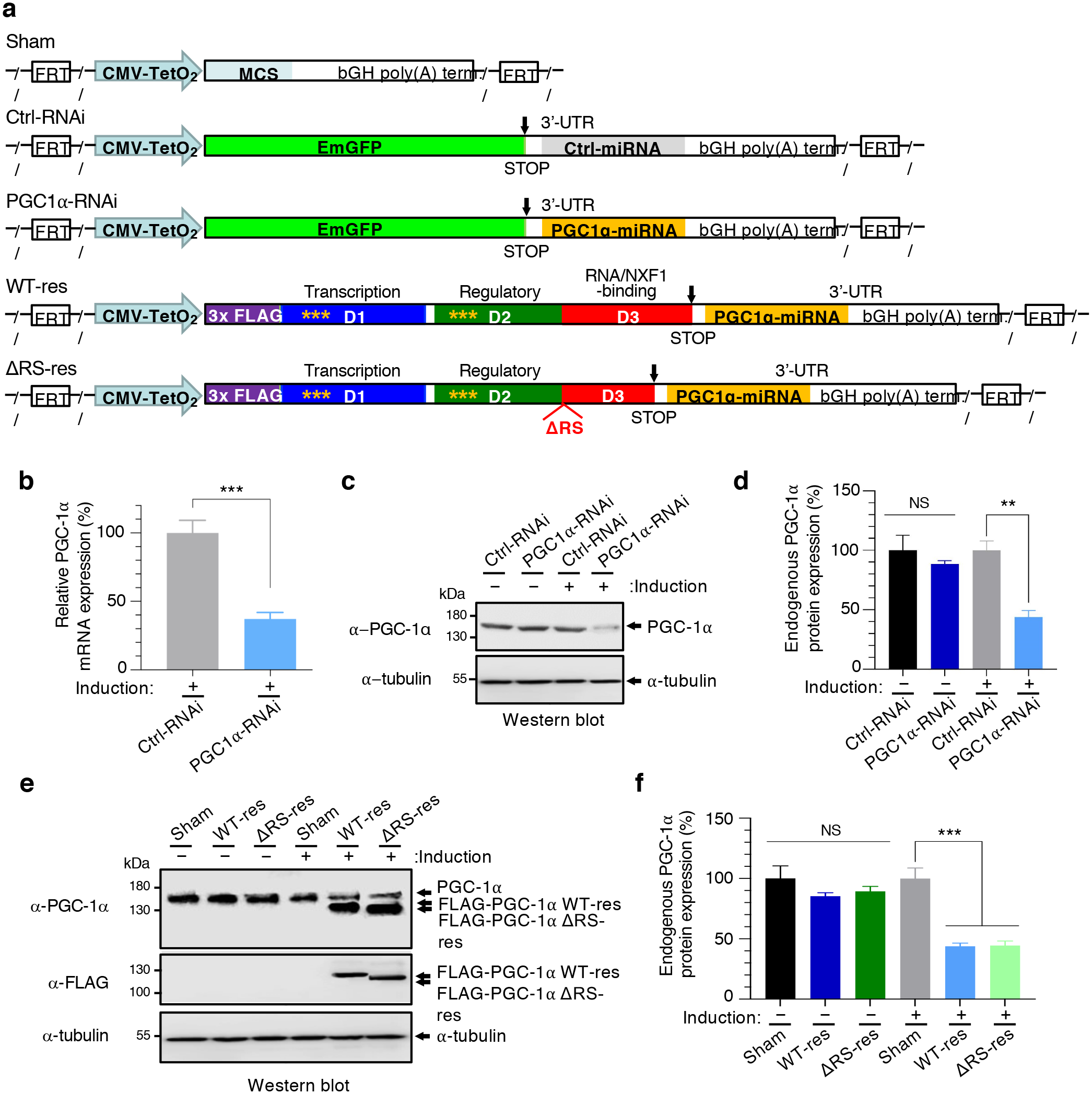
Engineering and testing the functionality of human isogenic stable cell lines. **(a)** Schematic representation of HEK293 Flp-In^TM^ T-REx^TM^ stable inducible cell line generated. Sham (Flp-In control) and either Ctrl-RNAi (scrambled non-targeting sequence) or PGC-1α RNAi stable cell lines were generated along with miRNA resistant N-terminally FLAG-tagged full length (WT) and PGC-1α-ΔRS cell lines which harbour the PGC-1α miRNA cassette in the 3’UTR. Yellow * highlight the position of the silent nucleotide changes which confer resistance to the miRNAs targeting endogenous PGC-1α transcripts. FRT indicates the plasmidic Flp Recognition Target sites which allows integration of the various transgenes at the single HEK293 Flp-In T-Rex genomic FRT site. **(b)** Endogenous PGC-1α transcript levels from of Ctrl-RNAi and PGC-1α RNAi cell lines induced with doxycycline for 72 h were quantified by qRT-PCR analysis following normalisation to U1 snRNA levels in three biological replicate experiments (Mean ± SEM; one-way ANOVA with Tukey’s correction for multiple comparisons; ***: p<0.001). **(c)** Western blots from Ctrl-RNAi and PGC-1α RNAi cell lines either with or without doxycycline induction for 72 h. Blots were probed for PGC-1α and α-Tubulin. **(d)** Western blots shown in (c) were quantified in triplicate experiments (Mean ± SEM; one-way ANOVA with Tukey’s correction for multiple comparisons; NS: not significant, **: p<0.01). **(e)** Western blots from Sham, PGC-1α WT-res and PGC-1α PGC-1α ΔRS-res cell lines either with or without doxycycline induction for 72 h. Blots were probed for PGC-1α, FLAG, which detects the induced miRNA resistant PGC-1α and α-Tubulin. **(f)** Western blots shown in (e) were quantified in triplicate experiments (Mean ± SEM; one-way ANOVA with Tukey’s correction for multiple comparisons; NS: not significant, ***: p<0.001).

### PGC-1α drives the nuclear export of transcriptionally co-activated mRNAs encoding mitochondria-related proteins

PGC-1α is known to co-activate the nuclear transcription of *TFAM*^5^, *NRF-1*^5^ and transcripts encoding cytochrome c oxidase (COX) subunits of the terminal mitochondrial respiratory chain complex IV, including COX assembly homolog 10 (*COX10*) ^59^ and COX subunit 5A (*COX5A*) ^60^. Formaldehyde cross-linked RNA immunoprecipitation (RIP) assays in induced human cell lines show that transcripts of known genes co-activated by PGC-1α specifically interact with FLAG-tagged PGC-1α but not with the ΔRS mutant deficient for RNA-binding (**Fig. 5a**). On the other hand, catalase (*CAT*), superoxide dismutase 2 (*SOD2*) and glyceraldehyde 3-phosphate dehydrogenase (*GAPDH*) mRNAs which are expressed from PGC-1α-independent promoters^59, 60^ do not interact with PGC-1α (**Fig. 5a**), indicating that PGC-1α binds these cellular mRNAs it transcriptionally co-activates. To functionally characterise the potential role of PGC-1α in the nuclear export of mRNAs, these transcripts were further quantified by qRT-PCR in total, nuclear and cytoplasmic fractions isolated from the doxycycline-induced cell lines. The quality of subcellular fractionation was validated using western blots probed by the nuclear remodelling factor SSRP1 and the cytoplasmic β-tubulin marker TUJ1, showing absence of nuclear contamination in the cytoplasmic fractionation (**Fig. 5b-c**). Doxycycline-induced depletion of PGC-1α leads to nuclear accumulation and concomitant cytoplasmic decrease of *TFAM*, *NRF1*, *COX10* and *COX5A* while the total levels of these transcripts remain unaffected in medium containing galactose (**Fig. 5d**), which forces cells to rely on mitochondrial metabolism through oxidative phosphorylation instead of glycolysis. The depletion of PGC-1α was validated using 2 pairs of primers annealing in either the coding sequences (CDS) or the 3’-untranslated region (3’-UTR) of the endogenous PGC-1α mRNA. Same results showed that PGC-1α also plays a role in the nuclear export of these transcripts in standard medium containing glucose (**Supplementary Fig. 5a-b**). In contrast, the expression levels of *SOD2*, *CAT* and *GAPDH*, transcripts which are not regulated^59, 60^ or bound (Fig. 5a) by PGC-1α, are not altered upon depletion of PGC-1α (Fig. 5d, Supplementary Fig. 5b). These assays demonstrate that PGC-1α licenses the nuclear export of the tested mRNAs it transcriptionally co-activates. Moreover, doxycycline-induced depletion of endogenous PGC-1α and expression of RNAi-resistant FLAG-tagged PGC-1α WT or ΔRS led to nuclear accumulation and cytoplasmic decrease of the *TFAM*, *NRF1*, *COX10* and *COX5A* mRNAs in the ΔRS-res cell line, showing that expression of the mutant PGC-1α lacking the RNA/NXF1-binding domain inhibits the nuclear export of PGC-1α target transcripts, while restoring the expression of the wildtype protein rescues the defects (**Fig. 5e**). In order to assess the functional relevance of the PGC-1α-dependent mRNA nuclear export function, we examined the expression levels of the TFAM protein and found that it has reduced expression levels in the induced PGC1α-RNAi and ΔRS-res cell lines, but not in control and rescue conditions (**Fig. 5f**; quantification in **Fig. 5g-h**). Taken together, our data demonstrate that the RNA-binding domain of PGC-1α exhibits a novel NXF1-dependent mRNA nuclear export function which is linked to the transcripts it co-activates.

**Figure 5.**
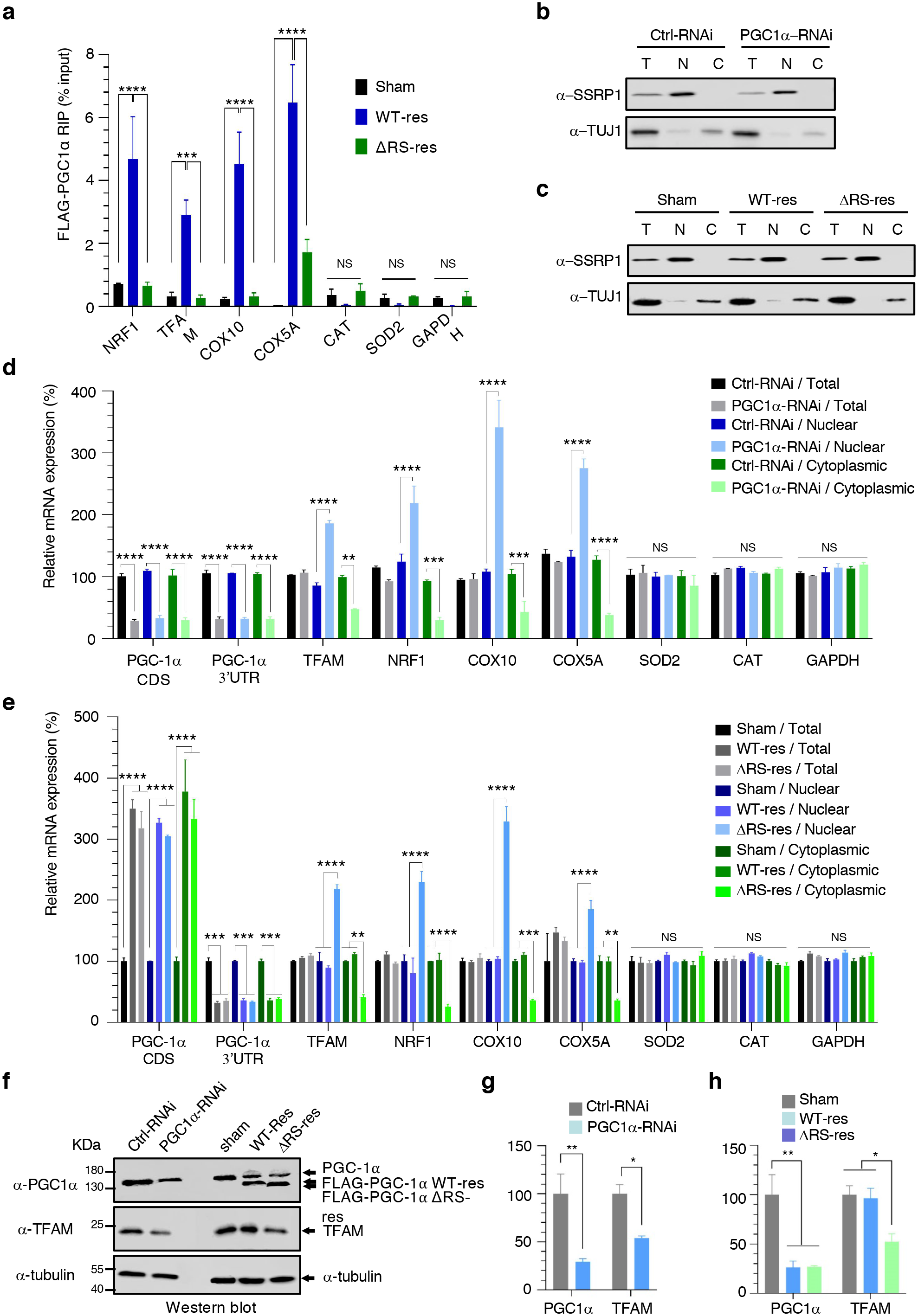
The RS domain of PGC-1α drives the nuclear export of co-activated mRNAs encoding mitochondrial-related proteins. (**a**) RNA immunoprecipitation (RIP) assay. Formaldehyde was added to Sham, FLAG-tagged RNAi-resistant PGC-1α WT (WT-res) and FLAG-tagged PGC1α-ΔRS (ΔRS-res) human stable cell lines induced for 6 days with doxycycline and subjected to anti-FLAG immunoprecipitation. Purified RNA was analysed by qRT-PCR and expressed as a percentage of the input. RIPs were performed in three biological replicate experiments (Mean ± SEM; two-way ANOVA with Turkey’s correction for multiple comparisons; NS: not significant, ***: p<0.001, ****: p<0.0001). **(b)** Western blots of Ctrl-RNAi and PGC1α-RNAi cell lines induced for 6 days with doxycycline and subjected to cellular fractionation using hypotonic lysis to yield cytoplasmic fractions. The chromatin remodelling SSRP1 factor is used to check for potential nuclear contamination in cytoplasmic fractions. Depletion of TUJ1 (beta-Tubulin III) in nuclear fractions was used to check for quality of the nuclear fractions. **(c)** Western blots of Sham, WT-res and ΔRS-res cell lines induced for 6 days with doxycycline and subjected to cellular fractionation using hypotonic lysis to yield cytoplasmic fractions. The chromatin remodelling SSRP1 factor is used to check for potential nuclear contamination in cytoplasmic fractions. Depletion of TUJ1 in nuclear fractions was used to check for quality of the nuclear fractions. **(d)** Total, nuclear and cytoplasmic levels of fractionated RNA transcripts from (b) were quantified in three biological replicate experiments by qRT-PCR following normalisation to U1 snRNA levels and to 100% in Ctrl-RNAi cell line (Mean ± SEM; two-way ANOVA with Tukey’s correction for multiple comparisons, NS: not significant, **: p<0.01, ***: p<0.001, ****: p<0.0001). **(e)** Total, nuclear and cytoplasmic levels of fractionated RNA transcripts from (c) were quantified in three biological replicate experiments by qRT-PCR following normalisation to U1 snRNA levels and to 100% in Ctrl-RNAi cell line (Mean ± SEM; two-way ANOVA with Tukey’s correction for multiple comparisons, NS: not significant, **: p<0.01, ***: p<0.001, ****: p<0.0001). The depletion of endogenous PGC-1α and expression of RNAi-resistant FLAG-tagged PGC-1α WT or ΔRS is validated with primers annealing in the 3’UTR of the PGC-1α gene or in the coding region of the transgene respectively. **(f)** Western blots from Ctrl-RNAi, PGC1α-RNAi, Sham, WT-res and ΔRS-res cell lines induced with doxycycline for 6 days. Blots were probed for PGC-1α, TFAM and α-Tubulin. **(g-h)** Western blots shown in (f) were quantified in three biological replicate experiments (Mean ± SEM; one-way ANOVA with Tukey’s correction for multiple comparisons; *: p<0.05, **: p<0.01).

### The mRNA nuclear export activity of PGC-1α is essential to its cellular function and mitochondrial homeostasis

*TFAM* and *NRF1* encode factors involved in stimulating the expression of genes involved in mitochondrial biogenesis, while *COX10* and *COX5A* transcripts encode subunits of the respiratory chain complex IV which are of critical importance for the assembly of the complex^61^. Western blots probed with OXPHOS antibodies, a cocktail of 5 antibodies for simultaneous detection of a representative protein in each of the mitochondrial complexes I to V, indicated that the expression level of mitochondrial-encoded complex IV COX2 is down-regulated upon depletion of PGC-1α (**Fig. 6a**) and expression of the RNA/NXF1-binding mutant (**Fig. 6b**). We further confirmed the expected galactose-mediated induction of the COX2 protein and the significant alteration of its expression levels in PGC1α-RNAi or ΔRS-res cell lines in galactose (**Fig. 6 c-d**) in agreement with the role of PGC-1α in the nuclear export of transcripts encoding proteins involved in mitochondrial biogenesis and complex IV assembly. On the other hand, the expression of proteins involved in complex I, III and V was not affected while, albeit not statistically significant, there was an upregulation of the SDHB protein composing complex II (**Supplementary Fig. 6a-b**).

**Figure 6.**
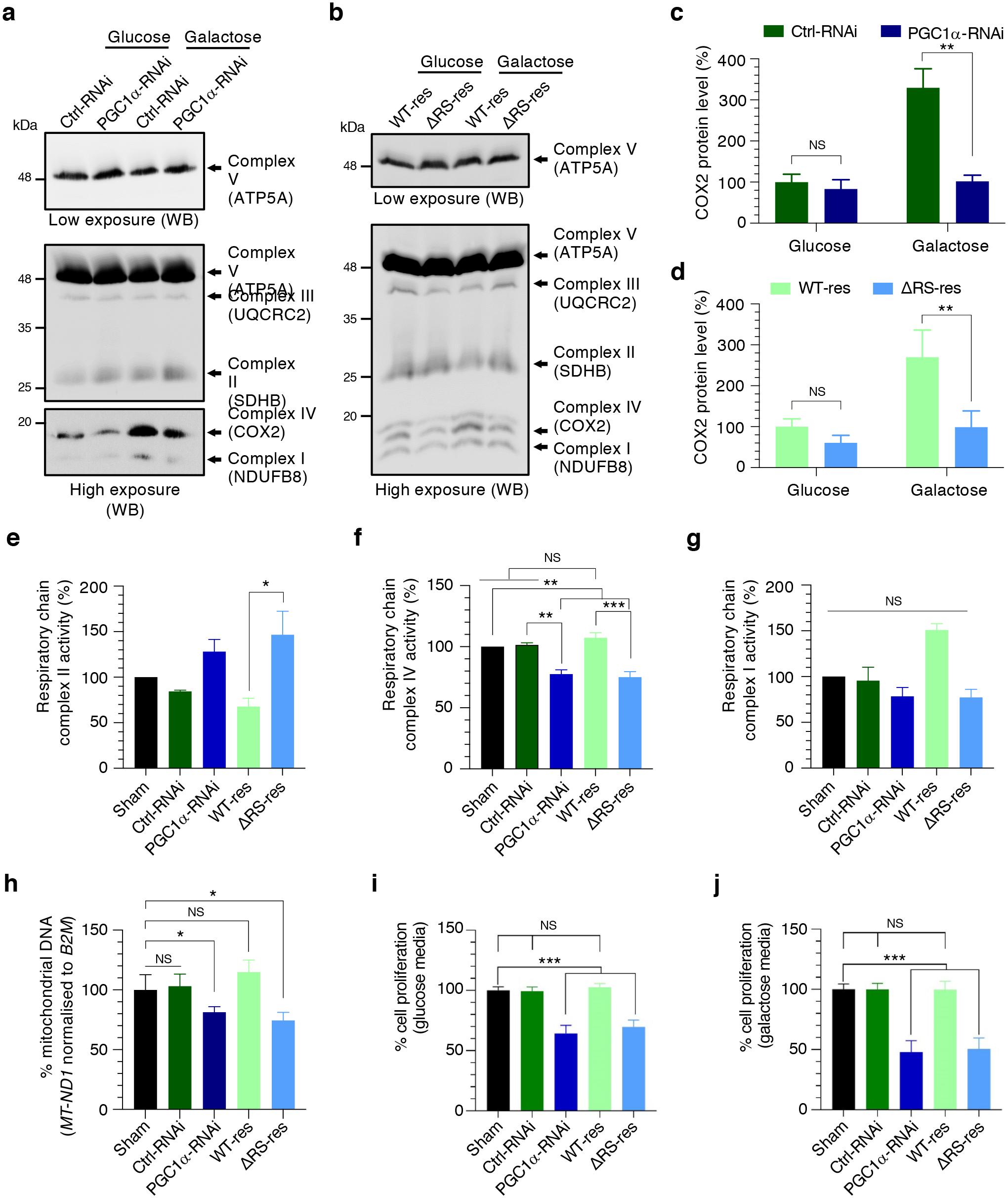
The mRNA nuclear export function of PGC-1α is essential to its cellular function and mitochondrial homeostasis. (**a**) OXPHOS Western Blots of complex I, II, III & IV protein levels of Ctrl-RNAi and PGC-1α RNAi cell lines induced for 72 h with doxycycline and cultured in either glucose or galactose media for the last 24 h prior to analysis. **(b)** OXPHOS Western Blots of complex I, II, III & IV protein levels of PGC-1α WT and PGC-1α ΔRS cell lines induced for 72 h with doxycycline and cultured in either glucose or galactose media for 24 h prior to analysis. **(c)** OXPHOS western blots shown in (a) were performed on three biological replicate experiments and changes in COX2 levels quantified and normalised against the total mitochondrial protein level (the sum of all 5 complexes) (Mean ± SEM; two-way ANOVA with Tukey’s correction for multiple comparisons; NS: not significant, **: p<0.01). **(d)** OXPHOS western blots shown in (b) were performed on three biological replicate experiments and changes in COX2 levels quantified and normalised against the total mitochondrial protein level (the sum of all 5 complexes) (Mean ± SEM; two-way ANOVA with Tukey’s correction for multiple comparisons; NS: not significant, **: p<0.01). **(e)** Mitochondrial **r**espiratory chain complex II enzymatic activity of Sham, Ctrl-RNAi, PGC1α-RNAi, WT-res and ΔRS-res cell lines induced for 6 days with doxycycline and cultured in galactose media for the last 24 h prior to analysis were performed in three biological replicate experiments (Mean ± SEM; one-way ANOVA with Turkey’s correction for multiple comparisons; *: p<0.05). **(f)** Mitochondrial **r**espiratory chain complex IV enzymatic activity of Sham, Ctrl-RNAi, PGC1α-RNAi, WT-res and ΔRS-res cell lines induced for 6 days with doxycycline and cultured in galactose media for the last 24 h prior to analysis were performed in three biological replicate experiments (Mean ± SEM; one-way ANOVA with Turkey’s correction for multiple comparisons; NS: not significant, **: p<0.01, ***: p<0.001). **(g)** Mitochondrial **r**espiratory chain complex I enzymatic activity of Sham, Ctrl-RNAi, PGC1α-RNAi, WT-res and ΔRS-res cell lines induced for 6 days with doxycycline and cultured in galactose media for the last 24 h prior to analysis were performed in three biological replicate experiments (Mean ± SEM; one-way ANOVA with Turkey’s correction for multiple comparisons; NS: not significant). **(h)** Genomic mitochondrial *MT-ND1* DNA levels were quantified from Sham, Ctrl-RNAi, PGC1α-RNAi, WT-res and ΔRS-res cell lines induced for 6 days with doxycycline and cultured in galactose media for the last 24 h prior to analysis. 5 biological replicate experiments were performed and analysed by qPCR following normalisation to the nuclear *B2M* gene and to 100% of Sham cell line (Mean ± SEM; one-way ANOVA with Tukey’s correction for multiple comparisons; NS: not significant, *: p<0.05). **(i)** MTT cell proliferation assay performed on Sham, Ctrl-RNAi, PGC1α-RNAi, WT-res and ΔRS-res cell lines induced for 6 days with doxycycline and cultured in glucose media were performed in 5 biological replicate experiments (Mean ± SEM; one-way ANOVA with Tukey’s correction for multiple comparisons; NS: not significant, ***: p<0.001). **(j)** MTT cell proliferation assay performed on Sham, Ctrl-RNAi, PGC1α-RNAi, WT-res and ΔRS-res cell lines induced for 6 days with doxycycline and cultured in galactose media prior to analysis were performed in 5 biological replicate experiments (Mean ± SEM; one-way ANOVA with Tukey’s correction for multiple comparisons; NS: not significant, ***: p<0.001).

We next sought to use more sensitive assays to measure the enzymatic activity of respiratory chain complexes I, II and IV in the induced WT-res and ΔRS-res cell lines. This confirmed that depletion of PGC-1α and specific alteration of its RNA/NXF1-binding function leads to increased activity of complex II (**Fig. 6e**) and inhibition of complex IV activity (**Fig. 6f**), providing functional validation of protein changes observed above. On the other hand, the activity of complex I is not statistically altered compared to the Sham control line (**Fig. 6g**). We also sought to test if the altered nuclear export of *TFAM* and *NRF1* mRNAs affect mitochondrial genome replication in the induced PGC1α-RNAi and ΔRS-res cell lines using qPCR analysis using an established protocol^62–64^. Expression of the unique and rarely deleted mitochondrial *MT*-*ND1* gene fragment is reduced upon depletion of PGC-1α and expression of the ΔRS mutant (**Fig. 6h**), consistent with a role of PGC-1α in the nuclear export of transcripts which encode proteins stimulating the biogenesis of mitochondria. Finally, to evaluate the physiological relevance of the RNA/NXF1-binding function of PGC-1α, we measured cell proliferation using MTT assays in Sham, Ctrl-RNAi, PGC1α-RNAi, WT-res and ΔRS-res cell lines. Both PGC1α-RNAi and ΔRS-res cell lines showed reduced cell proliferation with a more pronounced defect in galactose condition (**Fig. 6i-j**), a result consistent with dependence on mitochondrial respiration and increased requirement of PGC-1α activity. Taken together, our data demonstrate that the NXF1-dependent mRNA nuclear export function of PGC-1α is essential to mitochondrial metabolism homeostasis and optimal cellular growth.

### Proteome-wide implication of the mRNA nuclear export activity of PGC-1α

To determine the role of the RNA/NXF1-binding function of PGC-1α on the proteome, we analysed WT-res and ΔRS-res cell lines using 10plex Tandem Mass Tag (TMT) spectrometry, a methodology for multiplexed quantification of proteins in 10 samples (**Fig. 7a**). Principal component analysis of the identified proteomes validated the reproducibility of data in replicate samples (**Supplementary Fig. 7a**). Comparing WT-res and ΔRS-res proteomes of cells induced with doxycycline for 6 days indicated that, upon loss of the mRNA nuclear export function of PGC-1α, the expression levels of 216 and 124 proteins are respectively altered in glucose and galactose media (**Supplementary table 1 tabs 1-2**). Among these, 115 proteins (approximately 50%) were commonly altered in both glucose and galactose (**Supplementary Fig. 7b**), showing same direction of changes between glucose and galactose conditions (**Supplementary Table 1 tab 3**). To increase statistical power, and given that same directions of changes were identified for commonly altered hits in glucose and galactose, the differential expression analysis was performed again by combining the proteomes obtained from glucose and galactose (**Supplementary Table 1 tab 4**). It was also represented as a volcano plot showing down and up regulation of 209 and 130 proteins respectively (**Fig. 7b**). In particular subunits of the respiratory chain complex II, SDHA, SDHB and SDHAF2/4, were upregulated. A summary of all datasets is shown **(Supplementary Table 1 tab 5).** Gene ontology (GO) analysis was performed on the 339 proteins altered upon loss of the mRNA nuclear export function of PGC-1α (glucose + galactose analysis (Supplementary Table tab 4). This highlighted involvement of this RNA-processing activity in (i) biological processes involving “respiratory electron transport” and pathways involved in mitochondria-related pathways such as “succinate metabolic process”, “oxidation-reduction process”, “tricarboxylic acid cycle”, “ATP biosynthetic process” and “fatty acid β-oxidation” as well as (ii) “RNA transport”, “metabolic pathways” and “carbon metabolism” in the KEGG Pathway investigation (**Fig. 7c**). Mitochondrion further topped up the enrichment list in the GO analysis of cellular components (**Fig. 7c**). Strikingly, these data confirm the key functional importance of the mRNA nuclear export activity of PGC-1α in the mitochondrial metabolism homeostasis at proteome-wide level.

**Figure 7.**
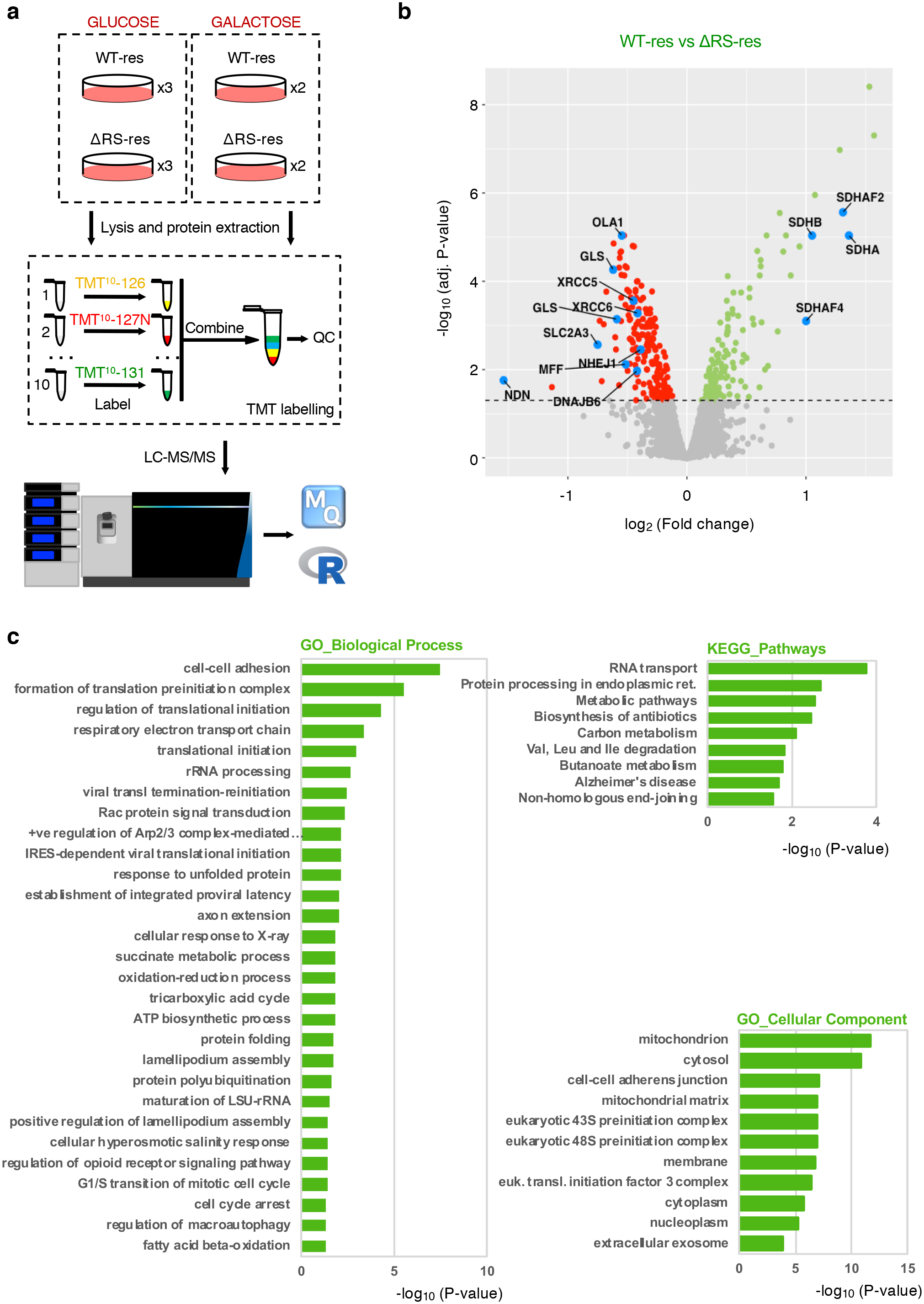
Proteome-dependence of the mRNA nuclear export function of PGC-1α. **(a)** Schematic of proteomics approach using Tandem Mass Tag (TMT) spectrometry to label samples for accurate multiplexing quantification of proteins (Methods). (**b**) Volcano plot showing log_2_ fold change (FC) of altered proteins identified in the proteomes of WT-res versus ΔRS-res cell lines induced with doxycycline for 6 days in media containing either glucose or galactose for the last 24 hours. The data show the combined proteomes identified in glucose and galactose media. Significantly down-(209, red label) and up-(130, green label) regulated proteins are highlighted for adjusted p-values <0.05 (Methods). **(c)** Gene ontology (GO) analysis of differentially-expressed proteins in the PGC-1α ΔRS cell line. Enrichment for biological processes, cellular components and KEGG Pathways are shown.

Interestingly, this investigation also highlighted reduced expression levels of proteins involved in processes not previously known to be regulated by PGC-1α. These include DnaJ Heat Shock Protein Family (Hsp40) Member B6 (DNAJB6) and Solute Carrier Family 2 Member 3 (SLC2A3, also known as GLUT3) (**Fig. 7b**, **Supplementary Fig. 7c**, **Supplementary Table 1 tab 5**) which were linked to Huntington’s disease^65, 66^, a disorder known to exhibit altered PGC-1α function. On the other hand, X-ray repair cross-complementing proteins 5 and 6 (XRCC5, XRCC6) and nonhomologous end joining factor 1 (NHEJ1) were also down-regulated upon loss of the mRNA nuclear export function of PGC-1α (**Fig. 7b**, **Supplementary Fig. 7c, Supplementary Table 1 tab 5**). Interestingly, they are linked to age-related telomere maintenance^67–69^, premature ageing and alteration of telomerase gene expression^70^. To functionally validate these findings, we tested whether these transcripts are novel gene expression targets regulated by PGC-1α. We also included telomeric repeat binding factor 2 (TERF2), a protein directly associated with age-related maintenance of telomeres^69, 70^. RNA immunoprecipitation assays in induced Sham, WT-res and ΔRS-res cells further showed that PGC-1α specifically binds *XRCC5*, *XRCC6*, *NHEJ1* and *TERF2* mRNAs via the RS domain (**Fig. 8a**). Moreover, while total mRNA expression levels are unchanged, the depletion of PGC-1α and/or expression of the RNA/NXF1-binding mutant lead to concomitant nuclear accumulation and cytoplasmic decrease (**Fig. 8b-c**), demonstrating that the RS domain of PGC-1α is involved in the nuclear export of these transcripts. These results show that the mRNA nuclear export activity of PGC-1α controls the gene expression of novel age-related PGC-1α-dependent target genes that were identified in our proteomics screen. They provide new mechanistic insights to previous studies reporting that increased and decreased level/activity of PGC-1α are associated with longevity and pre-mature ageing respectively^71, 72^, as well as with cell survival during stress^73^, linking thus mitochondrial function and lifespan to stress-resistance.

**Figure 8.**
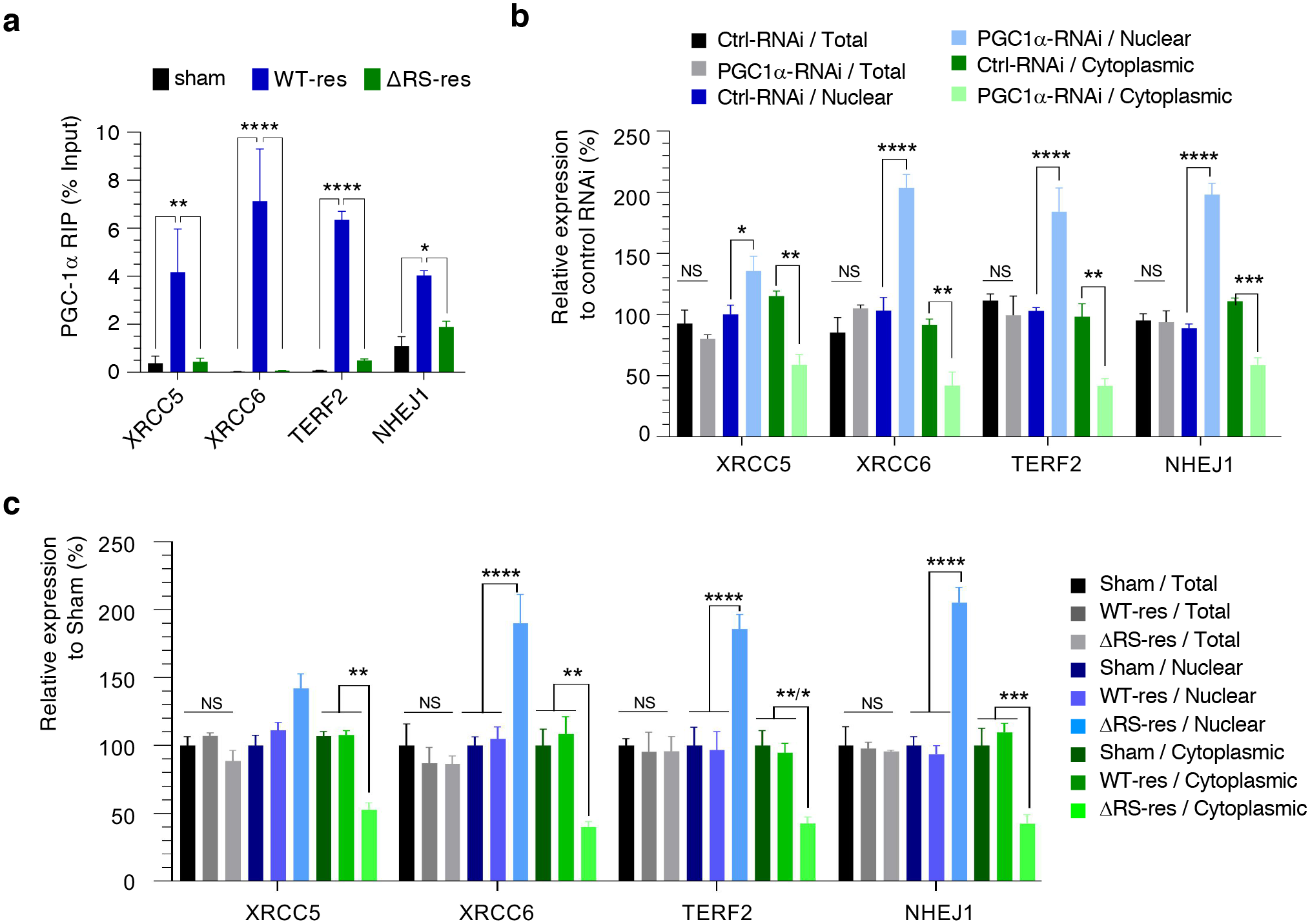
PGC-1α controls the nuclear export of transcripts encoding proteins involved in age-related telomere maintenance. (**a**) RNA immunoprecipitation (RIP) assay of transcripts down regulated in the TMT spectrometry analysis. Formaldehyde was added to Sham, FLAG-tagged PGC-1α WT and FLAG-tagged PGC-1α ΔRS stable cell lines induced for 6 days with doxycycline and subjected to anti-FLAG immunoprecipitation. Purified RNA was analysed by qRT-PCR and expressed as % of the input. RIPs were performed in three biological replicate experiments (Mean ± SEM; two-way ANOVA with Turkey’s correction for multiple comparisons; *: p<0.05, **: p<0.01, ****: p<0.0001). **(b)** Total, nuclear and cytoplasmic levels of fractionated RNA transcripts down regulated in the TMT spectrometry analysis were quantified in three biological replicate experiments by qRT-PCR following normalization to U1 snRNA levels and to 100% in ctrl-RNAi cell line (Mean ± SEM; two-way ANOVA with Tukey’s correction for multiple comparisons, NS: not significant, *: p<0.05, **: p<0.01***: p<0.001, ****: p<0.0001). **(c)** Total, nuclear and cytoplasmic levels of fractionated RNA transcripts down regulated in the TMT spectrometry analysis were quantified in three biological replicate experiments by qRT–PCR following normalization to U1 snRNA levels and to 100% in ctrl-RNAi cell line (Mean ± SEM; two-way ANOVA with Tukey’s correction for multiple comparisons, NS: not significant, *: p<0.05, **: p<0.01, ***: p<0.001, ****: p<0.0001).

## Discussion

Here, we characterised the RNA-binding function of the human master energy homeostasis regulator PGC-1α, showing that its RS domain interacts with RNA as well as the mRNA export receptor NXF1 to drive the nuclear export of mRNAs it transcriptionally co-activates.

Functional assays and proteomics showed that this novel mRNA nuclear export activity is essential for PGC-1α’s function in mitochondrial metabolism homeostasis and cell growth, its key physiological role. Interestingly, we also identified that the mRNA nuclear export function of PGC-1α is involved in novel gene expression targets related to age-associated maintenance of telomeres, a pathway directly relevant to its known roles in the control of premature ageing and longevity. The proteome studies increased the repertoire of potential new PGC-1α targets, highlighting that the expression levels of additional proteins in the energy metabolism, stress response or translation initiation are regulated by the mRNA nuclear export function. Additional studies will be needed to validate further the functional cellular implications of these findings.

Interestingly, PGC-1α was recently reported to interact with a large proportion of intronic RNA sequences in pre-mRNAs^44^, thus placing PGC-1α in an ideal position to further promote gene expression through co-transcriptional recruitment to PGC-1-activated transcripts and coupling to mRNA nuclear export^74–76^. In human, the bulk nuclear export of mRNAs is primarily linked to the splicing-dependent recruitment of the TREX complex^37–39^, which contains the mRNA nuclear export adaptor Aly/REF (also known as THOC4 subunit), and provides a platform for the high RNA-affinity remodelling of NXF1^40, 41^. On the other hand, the SR-rich splicing factors SRSF1,3,7 were not found associated to human TREX^77^ however the co-transcriptional recruitment of their phosphorylated forms contribute to pre-mRNA splicing and regulation of alternative splicing. Upon completion of splicing, the dephosphorylated forms of SRSF1,3,7 further link splicing to mRNA nuclear export through interactions with NXF1^31, 32^ and high RNA affinity remodelling of NXF1^33, 34, 40^ which licenses the transport of the mRNA through the channel of the nucleopore. In the yeast *Saccharomyces cerevisiae*, which harbors a genome with a low proportion of spliced genes, the TREX complex predominantly couples transcription elongation to nuclear mRNA export^78, 79^.

Unlike SR-rich splicing factors^77^, we found that PGC-1α interacts with several subunits of the human TREX complex. This interaction appears to be physiologically relevant since altering the RNA-binding activity of PGC-1α leads to increased interactions with TREX subunits, suggesting stalling of the dynamic NXF1-remodeling process which requires RNA. Interestingly, it was also reported that the yeast TREX complex recruits SRSF-like Gbp2 and Hbr1 proteins during transcriptional elongation^80^. More research is now required to investigate whether PGC-1α plays a role in the splicing of cellular transcripts in order to dissect the molecular mechanisms which couple the transcriptional co-activation and mRNA nuclear export functions of PGC-1α. The mRNA nuclear export function of PGC-1α could be linked to transcriptional co-activation via (i) a potential role of PGC-1α in splicing or (ii) directly through transcriptional activation of a subset of mRNAs expressed from genes with PGC-1α-regulated promoters. Interestingly, the general mRNA nuclear export adaptor Aly/REF, a subunit of TREX, was initially characterised as ALY (Ally of AML-1 and LEF-1) and BEF (bZIP enhancing factor) in studies reporting that it stimulates transcriptional activation by enhancing the binding of some transcription factors to DNA^81, 82^. We present a model which shows the recognised and novel functions of PGC-1α (**Fig. 9**). The discovery of this novel biological function for a key cellular homeostasis regulator is anticipated to provide a stepping step to progress PGC-1α research into fields covering gene expression, metabolic disorders, ageing and neurodegenerative diseases.

**Figure 9.**
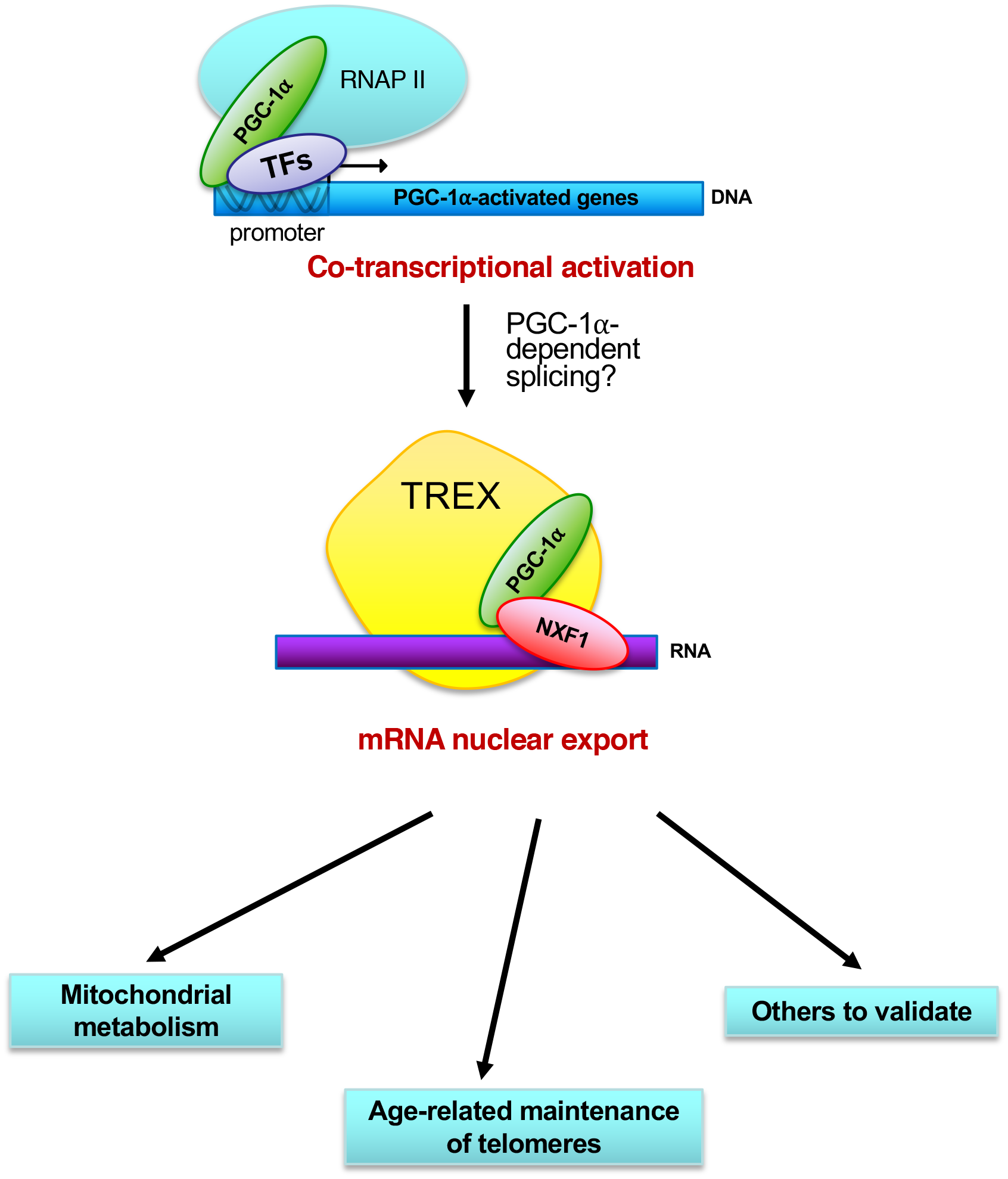
Model coupling the transcriptional co-activation and mRNA nuclear export functions of PGC-1α. The master regulator PGC-1α is well characterised for its role in binding and co-activating several transcription factors (TFs) to stimulate the RNA polymerase II (RNAPII) dependent transcription of target genes with PGC-1α-activated promoters which regulate the energy metabolism homeostasis. Here, we showed that PGC-1α binds transcripts it co-activates to further license their NXF1-dependent nuclear export into the cytoplasm, while its RNA-processing function is not associated with transcripts expressed from PGC-1α-independent promoters. It remains unknown whether an additional potential splicing function of PGC-1α may be involved in the coupling of the transcriptional co-activation and mRNA nuclear export functions. Interestingly, PGC-1α binds RNA via its RS domain in contrast to the SRSF1,3,7 factors which interacts semi-specifically with RNA via the RNA Recognition Motifs.

## Methods

### Plasmids

All plasmids used in this study are described in **Supplementary Table 2**. RNA interference plasmids were built using annealed DNA oligonucleotides (**Supplementary Note 1**) in the pcDNA6.2-GW/EmGFP plasmid (Block-iT™ miRNA expression kit from *Invitrogen*). Specific primers for pre-miRNA were designed using Life Technologies’ online tool (www.lifetechnologies.com/rnai). Single miRNAs were generated following manufacturer’s instructions (Block-iT; LifeTechnologies). Briefly, 50 µM of each top and bottom miRNA oligos were annealed together. Ten nM annealed oligos were used for molecular cloning. For the chaining, single miRNA was removed from one of the plasmids by double restriction digest and inserted into the other plasmid containing another miRNA. miRNAs were initially chained in pcDNA6.2-GW/EmGFP and the miRNA cassettes were subsequently amplified and cloned into the 3’UTR of an engineered pcDNA5-FRT/3xFLAG-PGC-1α plasmid.

### Generation and culture of HEK293T FlpIN isogenic stable inducible cell lines

Flp-In™ T-REx™ HEK293 parental cells (Invitrogen) were seeded at 1×10^6^ cells per 10 cm culturing dish in 15 ml media (DMEM High Glucose, 10% Tet-Free FBS (Sigma), 1% PenStrep) in the presence of selection antibiotic blasticidin S (Blast, 15 μg/ml, Calbiochem). Twenty-four h post plating, cells were co-transfected with FLP recombinase plasmid (pGKFLPobpA, Addgene) and gene of interest (GOI) (pcDNA5/FRT/3xFLAG-GOI) in a ratio 6:4. Forty-eight h post transfection, each confluent cell dish was split into 3x 10 cm dishes and cultured in 10 ml media mix (1:1 fresh:conditioned) in the absence of selection antibiotics. On the following day, another antibiotic hygromycin B (Hyg, 100 μg/ml, Invitrogen) was introduced into culture medium with Blast, and cells were cultured in Hyg/Blast selection medium for 7 days. The parental cell line carries dual resistance to blasticidin (near tetracycline repressor site) and zeocin (Zeo, 100 μg/ml, Invitrogen) at FRT site. During recombination, the zeocin resistance gene is replaced by hygromycin resistance gene which is acquired from the integration of pcDNA5/FRT-GOI plasmids. Cells that have failed to incorporate GOI at FRT site start to die during the culture in the selection medium containing hygromycin. At day seven, media was fully replaced with “fresh” media mix supplemented with Hyg/Blast and cells were cultured further for a week. During this period single colonies of cells carrying the GOI start to appear. Individual surviving colonies comprising of over 100 cells were then transferred to a single well on a 24-well plate and allowed to divide to confluency. At confluency, three-fifths of each well was transferred to a 6-well plate and cultured in media supplemented with Hyg/Blast. One-fifth of the cells were left in the same well and further cultured in Hyg/Blast selection medium while the remaining one-fifth was transferred to a neighbouring well and cultured in Zeo/Blast selection medium. Successful GOI incorporation was established when colonies survived in the media supplemented with Hyg/Blast but died in the presence of Zeo. Successful clones were further expanded to 10 cm dishes before being frozen down and stored in liquid nitrogen. Individual stable inducible cell lines were selected for downstream experiments after functional characterisation of the cells. The selection criteria were their ability to express GOI after doxycycline (1 μg/ml) induction for 72 h, normal doubling growth rate between 24-36 h, and healthy HEK293-like morphology.

For all experiments, cells were seeded in T175 flasks (5×10^6^ cells), 10 cm dishes (2×10^6^ cells), 6-well plates (2×10^6^ cells) or 24-well plates (25,000 cells) in media plus Hyg/Blast. Cells were induced with 1 µg/ml doxycycline 24 h post plating. For 6 day inductions, cells were split 72 h after induction to maintain optimum confluency into fresh media plus Hyg /Blast and 1 µg/ml doxycycline. For galactose media experiments, cells were changed into galactose media 24 h prior to assaying using Dulbecco’s Modified Eagles Medium containing no glucose (Gibco™) supplemented with 10% TET-free FBS, 5 U/ml Penstrep and 5 mM galactose.

### Culture and transfection of HEK 293T cells

HEK293T cells were seeded in 10 cm dishes (2×10^6^ cells) in media (DMEM High Glucose, 10% FBS (Sigma), 1% PenStrep) Twenty-four h post plating, cells were transfected with 3.5 µg polyethylenimine (PEI) / ml media. 15 μg or 500-700 ng total plasmid DNA were respectively transfected in 10 cm or in each well of a 24-well plate.

### Recombinant protein expression and purification of GB1-tagged PGC-1α domains

All PGC1α domains were cloned into pET24b-GB1-6xHis vector and transformed into *E. coli* BL21-RP cells. The expression of recombinant PGC-1α proteins was induced for 3 h at 37°C with 0.4 mM isopropyl-β-D-thiogalactoside (IPTG, Calbiochem) in Terrific Broth and purified by metal ion affinity chromatography (IMAC) on TALON/Cobalt beads (Clontech) using 1M NaCl containing buffers to prevent the potential co-purification of *E. coli* RNA (Lysis buffer: 50 mM Tris-HCl pH8.0, 1M NaCl, 0.5% Triton X-100; Wash buffer: 50 mM Tris-HCl pH8.0, 1M NaCl, 0.5% Triton X-100, 5 mM imidazole). Elution was achieved with the wash buffer containing 200 mM imidazole (50 mM Tris-HCl pH8.0, 500 mM NaCl, 200 mM imidazole). Domain 1 of PGC1α was readily soluble, however, the rest of the domain proteins were all insoluble and present in inclusion bodies in the bacterial pellets (Supplementary Figure 1). These insoluble domain proteins were therefore solubilised and purified in denaturing conditions using buffers containing 8 M urea.

For the purification of insoluble domain proteins from insoluble bacterial pellets, they were resuspended in Urea Loading Buffer (50 mM Tris at pH 8, 500 mM NaCl, 0.5% v/v Triton X-100, 8M urea) and left shaking for 1 h at room temperature. Supernatants were collected after 15 min centrifugation at 17,000 *g* at room temperature and then loaded onto Urea Loading Buffer equilibrated cobalt beads for 1 h incubation at room temperature. After 4 times washes with Urea Wash Buffer (50 mM Tris at pH 8.0, 500 mM NaCl, 5 mM Imidazole, 0.5% v/v Triton X-100, 8M Urea), bound proteins were eluted with Urea Elution Buffer (50 mM Tris at pH 8, 500 mM NaCl, 200 mM Imidazole, 0.5% v/v Triton X-100, 8M Urea). All proteins purified in Urea denaturing condition were dialysed in 1X RNA Binding Buffer (15 mM HEPES at pH 7.5, 5 mM MgCl_2_, 0.05% v/v Tween 20, 10% v/v glycerol) supplied with 100 mM NaCl and 5 mM Arg/Glu and 1 mM EDTA. After dialysis, the soluble proteins were collected from supernatant after 1 min centrifuge at 17,000 *g* and protein concentration was quantified via Bradford assay.

### RNA-protein UV cross-linking assay

1.5 μg RNA probes (5xAAAAUU or 5xGGGGCC) (*Dharmacon*) were end-radiolabelled with *γ*[^32^P]-ATP (*PerkinElmer*) using T4 polynucleotide kinase (PNK, NEB). Reaction was set up by following the manufacturer’s instructions. For the protein-RNA binding assay, 600 ng purified recombinant proteins were incubated in 1x RNA Binding Buffer (15 mM HEPES pH 7.9, 100 mM NaCl, 5 mM MgCl_2_, 0.05% v/v Tween 20, 10% v/v glycerol)with ∼20 nmol radiolabelled RNA for 20 min at room temperature followed by 20 min on ice prior to UV-irradiation for 10 min at 1.5 J/cm^2^. The reaction was terminated with 4x Laemmli Sample Buffer and heating at 95°C for 5 min. The RNA-protein binding complex was resolved on SDS-PAGE prior to analysis by Coomassie staining and PhosphoImaging (Typhoon FLA 7000, GE Healthcare).

### mRNP capture assay

HEK293T cells were seeded in 10 cm dishes (2 per condition) and transfected with 3xFLAG-PGC-1α WT or *Δ*RS. Forty-eight h post transfection, 1 dish for each condition was UV-crosslinked on ice at 0.3 J/cm^2^ in 1 ml DEPC-treated PBS. All plates were then washed with ice cold DEPC PBS and lysed in 500 μl lysis buffer (50 mM Tris-HCl pH 7.5, 100 mM NaCl, 2 mM MgCl_2_, 1 mM EDTA, 0.5% v/v Igepal C9-630, 0.5% w/v Na-deoxycholate) supplemented with fresh Complete protease inhibitor cocktail (Roche) and 20U Ribosafe (Bioline) on ice for 5 min. Samples were centrifuged for 10 min at 17,000 *g* at 4°C. Cleared lysates were diluted to 2 mg/ml with lysis buffer, input samples kept, and diluted with equal volume 2x Binding Buffer (20 mM Tris-HCl pH7.5, 1M NaCl, 1% SDS, 0.2 mM EDTA). Mix was incubated with 25 μg oligo-dT beads rotating end-over-end at room temperature for 1 h. Beads were washed 3 times with 1x Binding Buffer and mRNP complexes were eluted in 50 μl Elution Buffer (10 mM Tris-HCl pH7.5, 1mM EDTA) supplemented with Complete protease inhibitor and 10 μg RNase A (Sigma) at 37°C for 30 min with gentle agitation. Inputs and eluates were subjected to western blotting.

### RNA immunoprecipitation (RIP) assays

HEK FlpIN cell lines Sham, PGC1α WT-res and PGC1α ΔRS-res were used in this study. Cells were cultured in a T175 flask and induced for 6 days. Cells were switched to galactose media 24 h prior to assaying. 1% formaldehyde was added to the medium of live cells for 10 minand subsequently quenched with 250 mM Glycine for 5 min at room temperature. Cells were washed with DEPC-treated PBS prior to being scraped into ice cold RNase-free IP150 lysis buffer (DEPC-treated water containing 50 mM HEPES pH 7.5, 150 mM NaCl, 10% glycerol, 0.5% Triton X-100, 1 mM EDTA, 1 mM DTT, 1 µl RNase inhibitor, protease inhibitors). Cells were passed through a 21G needle 10 times and left to lyse on ice for 10 min, followed by centrifugation at 17,000 *g* at 4°C for 5 min and quantification using Bradford Reagent. 2 mg of total protein at a 1 mg/ml concentration was incubated with 40 µl anti-FLAG-M2 affinity resin (Merck) (which had been blocked overnight with 1% BSA and 5 µl/ml ssDNA) overnight at 4°C on a rotating wheel. Beads were washed 5 times with RNase-free lysis buffer followed by elution with 50 µl RNase-free lysis buffer supplemented with 100 µg/ml 3xFLAG peptide (Sigma #F4799) for 30 min at 4°C on a rotating wheel. The formaldehyde crosslinks were reversed by heating the samples for 1 h at 70 °C and RNA was extracted using PureZOL™ (*BioRAD*) as described in the RNA extraction section. Extracted RNA samples were re-suspended in 25 µl (input) or 15 µl (eluate) RNase-free water.

### GST-NXF1:p15 pull-down assays

BL21-RP cells co-transformed with pGEX4T1/GST-NXF1 and pET9a/p15 were induced overnight at 18°C with 0.4 mM isopropyl-β-D-thiogalactoside (IPTG, Calbiochem) in Terrific Broth. Bacterial pellets (0.25 g) containing IPTG induced GST-tagged NXF1-p15 protein complex were lysed in RB-100 (25 mM HEPES pH 7.5, 100 mM potassium acetate, 10 mM MgCl_2_, 1 mM DTT, 10 % Glycerol and 0.5% Triton x-100), the soluble proteins were collected after a 5 min centrifuge at 17,000 *g* at 4 ℃ and immobilised on Glutathione-coated sepharose beads (*GE Healthcare*). For pull down with ^35^S-radiolabelled preoteins, PGC-1α full-length and protein domain were translated *in vitro* in the T7 transcription and translation coupled reticulocyte system (*Promega*) supplemented with ^35^S-Methionine. ^35^S-radiolabled proteins were incubated with the immobilized GST-NXF1-p15 protein complex in RB-100 buffer with or without RNaseA (20 ng/µl) at 4°C for 1 h. GST-NXF1:p15 bound protein complexes were eluted in GSH elution buffer (50 mM Tris, 100 mM NaCl, 40 mM reduced glutathione, pH 7.5) after 5x wash with binding buffer. The protein binding complex was separated on SDS-PAGE prior to analysis by Coomassie staining and Phosphoimaging (Typhoon FLA 7000, GE Healthcare). For pull down with purified PGC-1α protein domains, 25 µg purified recombinant proteins (see above) were added to the GST-NXF1:p15 binding reactions prior to incubation, wash and elution as highlighted for the pull down with the ^35^S-radiolabelled proteins. Co-purification of 6His-tagged PGC-1α protein domains was detected by western blot using anti-6His antibody (clone His.H8, ThermoFisher Scientific MA1-21315).

### Co-immunoprecipitation assays

HEK293T cells were seeded in 10 cm dishes and transfected with a 1:1 plasmid mixe of 13xMyc-tagged NXF1 and either 3xFLAG-GFP (1 dish) or 3xFLAG-PGC-1α plasmids (four dishes / construct). Forty-eight h post transfection, cells were washed with ice-cold PBS and lysed in 500 μl IP150 lysis buffer (50 mM HEPES pH 7.5, 100 mM NaCl, 1 mM EDTA, 1 mM DTT, 0.5% v/v Triton X-100, 10% v/v Glycerol) supplemented with protease inhibitors. Lysates were passed through a 21G needle ten times before centrifugation for 5 min at 17,000 *g* at 4°C. Cleared lysates were diluted to 1.5 mg/ml and incubated with 30 μl pre-blocked (2% BSA) anti-FLAG M2 affinity resin in the presence of 10 μg RNase A for 1.5 hr at 4°C rotating. Beads were then washed twice with lysis buffer and eluted with 100 μg/ml 3xFLAG peptide rotating for 1 h at room temperature. Protein complexes (co-immunoprecipitates) and input samples were subjected to western blotting.

### Confocal immunofluorescence microscopy

HEK293T cells were plated onto cover slips coated with 0.5 mg/ml gelatin solution (Sigma, G1393) in a 241--well plate. Forty-eight h later cells were washed with PBS and fixed with 4% paraformaldehyde for 10 min gently shaking at room temperature. Fixative was removed and wells were washed with PBS 3 times. Cells were blocked and permeabilised at the same time using blocking buffer (10% normal goat serum and 0.1% Triton X-100 in PBS) shaking for 30 min at room temperature. Cells were then incubated with primary antibodies (NXF1 ab129160 at 1:200; PGC-1α ST1202 at 1:1,000) in blocking buffer shaking at 4 °C overnight. Wells were washed with PBS 3 times of 10 min followed by a 2 h incubation at room temperature with secondary antibodies (rabbit AlexaFluor488 711-545-152 and mouse AlexaFluor647 715-605-151 at 1:500) in blocking buffer. Cells were washed 3 times with PBS where the second wash contained DAPI staining (dilactate, Invitrogen at 1:10,000). Coverslips were mounted onto glass slides using Fluoromount-G mounting medium (0100-01, SouthernBiotech) and left to dry. Slides were imaged with Leica SP8 Inverted confocal microscope using 63X oil immersion objective at 5X optical zoom. Images were acquired sequentially for each fluorescent channel as a Z-stack with a step size of 0.2-0.3 micron. For the analysis, confocal images were maximum intensity projected using FIJI software and analysed in ImageJ using the RGB Plot Plugin.

### Oligo-dT Fluorescent *In-situ* Hybridization (FISH)

HEK293T cells were seeded in 24-well plates (25,000 cells)and transfected with 3xFLAG-e1BAP5, 3xFLAG-Aly/REF or 3xFLAG-PGC-1α. Forty-eight h post transfection, cells were treated with 5 μg/ml Actinomycin D for 2 h followed by 3 washes with PBS and incubation in fix solution (4% PFA, 0.2% Triton X-100) for 30 min at room temperature. Wells were then washed 3 times with PBS and coverslips were transferred onto a Whatman paper in a petri dish. Each coverslip was incubated with 80 μl hybridization mix (20% Formamide, 2X SSC, 10% Dextran sulphate, 1% BSA, 4 μl ssDNA from salmon sperm, 1 μg Cy3-oligo-dT probe (Sigma)) for 2 h in the dark at 37°C. Coverslips were then washed 3 times with PBS and blocked with 2% BSA in PBS for 20 min at room temperature. Cells were stained with anti-3xFLAG antibody (Sigma) followed by AlexaFluor 488 (Abcam) secondary antibody and Hoechst stain. Mounted coverslips were imaged using Nikon LV100ND microscope with DS Ri1 Eclipse camera and images were processed using ImageJ software package.

### MTT cell proliferation assay

HEK FlpIN cell lines (Sham, Control-RNAi, PGC1α-RNAi, PGC1α WT-res and PGC1α ΔRS-res) were induced for 6 days with and plated into 24-well plates, cells were either maintained in glucose media or switched to galactose media 24 h prior to assaying.. Each plate contained 4 wells with only media to serve as a blank and 4 wells/cell line. For MTT assay, 250 mg Thiazolyl Blue Tetrazolum Bromide reagent (MTT) was added to each well and incubated in the dark at 37°C for 1 h. Cells were subsequently lysed with equal volume MTT lysis buffer (20% SDS, 50% Dimethylformamide (DMF)) and incubated, shaking, at room temperature for 1 h. Absorbance at 595 nm was then assessed with a PHERAstar FS (*BMG Labtech*). Absorbance data was retrieved using PHERAstar MARS (*BMG Labtech*), % cell proliferation calculated for each cell line relative to Sham and data analysed using GraphPad Prism (Version 9.0.2.161).

### Nuclear/ Cytoplasmic fractionation

HEK FLpIN cell lines (Sham, Control-RNAi, PGC1α-RNAi, PGC1α WT-res and PGC1α ΔRS-res) were induced for 6 days and cultured in 6-well plates (9 wells per cell line). Cells Cells were either maintained in glucose media or switched to galactose media 24 h prior to fractionation. Cytoplasmic fractionation was performed as described in^49^. Briefly, cells from 6 wells were collected in DEPC PBS and pelleted by centrifugation at 800 *g* for 5 min. Cell pellets were quickly washed with hypotonic lysis buffer (10 mM HEPES pH 7.9, 1.5 mM MgCl_2_, 10 mM KCl, 0.5 mM DTT) and lysed for 10 min on ice in hypotonic lysis buffer containing 0.16 U/µl RNase inhibitorand protease inhibitors. All lysates underwent differential centrifugation at 1,500 *g* (3 min), 3,500*g* (8 min), and 17,000*g* (1 min) at 4°C and the supernatants transferred to fresh tubes after each centrifugation. Nuclear pellets obtained after centrifugation at 1,500*g* for 3 min were lysed in Reporter lysis buffer (Promega), passed through a 21Gneedle and incubated on ice for 10 min before centrifugation at 17,000*g*, 5 min, 4 °C. Total fractions were collected from 3 wells of a 6-well plate in Reporter lysis buffer containing 16 U µl^−1^ RNase inhibitor protease inhibitorss and passed through a 21G needle prior to lysis for 10 min on ice before centrifugation at 17,000*g*, 5 min, 4°C. Equal volumes of total, nuclear and cytoplasmic lysates were subjected to western immunoblotting using SSRP1 and class III beta-tubulin antibodies. Total and fractionated extracts were added to PureZOL™ to extract the RNA.

### SDS-PAGE and Western Blotting

HEK FLpIN cell lines (Sham, Control-RNAi, PGC1α-RNAi, PGC1α WT-res and PGC1α ΔRS-res) were induced for 6 days and cultured in 6-well plates (1 well per cell line). Cells were washed in ice-cold PBS, scraped into ice-cold IP150 lysis buffer and left to lyse on ice for 10 min followed by centrifugation at 17,000 *g* at 4 °C for 5 minutes. Protein extracts were quantified using Bradford Reagent (BioRAD), resolved by SDS–PAGE, electroblotted onto nitrocellulose membrane and probed using the relevant primary antibody. PGC-1α (4C1.3) [1:1000 dilution] (Calbiochem ST1202), α-tubulin [1:10,000 dilution] (Sigma, clone DM1A), FLAG [1:2,000 dilution] (*Sigma* F1804, clone M2), ALYREF [1:2,000 dilution] (Sigma A9979, clone 11G5), TFAM (C-9) [1:1000] (Santa Cruz sc-376672), c-myc [1:1000] (Thermo Fisher MA1-980), 6xHIS [1:2000] (Proteintech 66005-1), NXF1 [1:2000] (Abcam ab50609), SSRP1 [1:1000 dilution] (Abcam ab26212) and OXPHOS [1:1000] (Abcam ab110411) antibodies were detected using HRP-conjugated mouse secondary antibody (Promega). UAP56/DDX39B [1:2000] (custom generated^51^) and THOC1 [1:1000] (Abcam ab131268) antibodies were detected using HRP-conjugated rabbit secondary antibody (Promega). Anti-Beta Tubulin III (Tuj1) [1:1000] (Millipore AB9354) antibody was detected using HRP-conjugated chicken secondary antibody (Promega).

### RNA extraction and qRT-PCR

PureZOL™ was added to lysates as per manufacturers recommendations and cleared by centrifugation for 10 min at 12,000 *g*, 4°C. Samples were incubated at room temperature for 10 min followed by the addition of 1/5^th^ volume of chloroform and vigorous shaking for 15 s. After a further 10 min incubation at room temperature, tubes were centrifuged at 12,000 *g* for 10 min (4 °C) and the upper phase collected. RNA was precipitated overnight at −20 °C with equal volume isopropanol, 1/10^th^ 3M NaOAc, pH5.8 and 1 µl Glycogen (5 µg µl^−1^, Ambion) and subsequently pelleted at 12,000 *g* for 20 min (4 °C). Pellets were washed with 70% DEPC ethanol and re-suspended in DEPC water. All extracted RNA samples were treated with DNaseI (Roche) and quantified using a Nanodrop (NanoDropTechnologies). Following quantification, 2 µg RNA (fractionation) or 11µl RNA (RNA immunoprecipitation) was converted to cDNA using BioScript Reverse Transcriptase (Bioline). Primers used in this study are provided in **Supplementary Table 3**. qPCR reactions were performed in duplicate using the Brilliant III Ultra-Fast SYBR Green QPCR Master Mix (Agilent Technologies) on a C1000 Touch™ thermo Cycler using the CFX96™ Real-Time System (BioRAD) using an initial denaturation step, 45 cycles of amplification (95°C for 30 s; 60°C for 30 s; 72°C for 1 min) prior to recording melting curves. qRT–PCR data was analysed using CFX Manager™ software (Version 3.1) (BioRAD) and GraphPad Prism (Version 9.0.2.161).

### Mitochondrial DNA quantification

HEK FLpIN cell lines (Sham, Control-RNAi, PGC1α-RNAi, PGC1α WT-res and PGC1α ΔRS-res) were induced for 6 days and cultured in 6-well plates (1 well per cell line). Cells were switched to galactose media 24 h prior to assaying. Genomic DNA was extracted using the DNeasy Blood and Tissue DNA extraction kit (Qiagen) as per the manufacturer’s protocol. Extracted DNA was quantified using a Nanodrop (NanoDropTechnologies). Mitochondrial DNA levels were assessed by qPCR using 100 ng genomic DNA and 300 nM primers (**Supplementary Table 3**) for the mitochondrial gene *MT*-*ND1* and the nuclear-encoded gene β-2 microglobulin *B2M*^63–64^. Reactions were performed in duplicate using the Brilliant III Ultra-Fast SYBR Green QPCR Master Mix (Agilent Technologies) on a C1000 Touch™ thermo Cycler using the CFX96™ Real-Time System (BioRAD). Amplification conditions were: an initial denaturation step of 95°C for 10 min followed by 40 cycles of amplification (95°C for 15 s; 60°C for 1 min; 72°C for 1 min) prior to recording melting curves. qRT–PCR data was analysed using CFX Manager™ software (Version 3.1) (BioRAD) and GraphPad Prism (Version 9.0.2.161).

### Activity of mitochondrial respiratory chain complexes I, II and IV

HEK FLpIN cell (Sham, Control-RNAi, PGC1α-RNAi, PGC1α WT-res and PGC1α ΔRS-res) were induced for 6 days and cultured in 10cm plates (2 plates per cell line). Cells were switched to galactose media 24 h prior to assaying. The activity of the mitochondrial complexes I, II and IV were measured using microplate activity assays (Abcam; ab109721, ab109908, ab10990). Assays were performed according to the manufacturer’s instructions using 20-35 µg protein per well. Activity reading were normalised to protein content. Sample protein concentrations were measured using Pierce BCA assay.

### Tandem Mass tag (TMT) spectrometry and data processing

HEK FLpIN cell (Sham, PGC1α WT-res and PGC1α ΔRS-res) were induced for 6 days and cultured in 6-well plates (1 well per cell line). Cells were either maintained in glucose media or switched to galactose media 24 h prior to assaying.were Cells washed with PBS containing protease inhibitors, gently scraped and flash frozen in liquid nitrogen followed by storage at −80°C. Three replicates were used for cells cultured in glucose, and two for those cultured in galactose giving a total of 10 samples. Each replicate represents a subsequent passage. Cell pellets were lysed in Urea/SDS lysis buffer (8 M Urea, 1% SDS, 50 mM HEPES pH 8.2, 10 mM Glycerol-2-phosphate, 50 mM NaF, 5 mM Sodium pyrophosphate, 1 mM EDTA, 1 mM Sodium vanadate, 1 mM DTT, 500 nM Okadaic acid, protease inhibitors, Phosphatase inhibitor cocktail III). After protein concentration was measured using Pierce BCA assay, 100 μg of each sample was used further for multiplexed quantitative proteomics giving a total of 1 mg protein. Each sample was reduced with 10 mM TCEP for 1 h at 55°C and alkylated with 20 mM IAA for 30 min in the dark at room temperature. In order to remove SDS, each reaction was precipitated with 6 volumes of cold (−20°C) acetone and left overnight at −20°C. Samples were equilibrated to room temperature to avoid urea crystals and centrifuged at 15,000 *g* for 10 min at room temperature. Supernatant was carefully removed and pellets were air-dried. Pellets were resuspended in 100 mM HEPES pH 8.5 by vortexing and incubation in a sonic bath for 15 min. Resuspended samples were digested with LysC (Lysyl endopeptidase, 125-05061, FUJIFILM Wako Chemicals) and Trypsin (MS grade, 90058, ThermoFisher Scientific) at 37°C shaking overnight. Each sample was then tandem mass tag (TMT) labelled for an hour at room temperature using a TMT10plex Isobaric Label Reagent Set (0.8 mg per tag, 90110, ThermoFisher Scientific; LOT UK282322) and following manufacturer’s instructions. A small aliquot of each sample was collected for a labelling efficiency and mixing accuracy quality checks (QCs) by liquid chromatography tandem mass spec (LC-MS/MS) using Orbitrap Fusion Lumos mass spectrometer and a 60 min HCD MS2 fragmentation method. The rest of the sample was stored at −80°C until QC results. Labelling efficiency of higher than 99% was obtained for each reaction and a mixing accuracy with lower than 1.5x difference between samples with lowest and highest summed intensity. Samples were defrosted, quenched with hydroxylamine for 15 min at room temperature and pooled together. Combined mixture was partially vacuum-dried and acidified to pH 2 followed by sample clean-up using C_18_ Sep Pak Vac 1cc, 50mg bed volume (Waters) with gravity flow. Final mixing check was performed by LC-MS/MS with a 240 min HCD MS2 fragmentation method. From the pooled labelled mix, 100 μg total (starting) protein was aliquoted and vacuum-dried to completion. Dried peptide mix was resuspended in 2% TFA and fractionated using Pierce High pH Reversed-Phase Fractionmation kit (84868, ThermoFisher Scientific). Each fraction was vacuum-dried and resuspended in 25 μl 0.1% TFA. Peptides were separated on a 50 cm, 75 μm I.D. Pepmap column over a 2 h gradient and eluted directly into the mass spectrometer (Orbitrap Fusion Lumos) with 3 injections of 7 μl for each fragmentation method – HCD MS2, and two methods for MS3^83, 84^. Xcalibur software was used to control the data acquisition. The instrument was run in data dependent acquisition mode with the most abundant peptides selected for MS/MS by HCD fragmentation.

Raw data were processed using MaxQuant v1.6.2.10 and Uniprot human reference proteome from September 2020. Processed data were then analysed using an R-coding script tailored to isobaric labelling mass spectrometry. The script was generated as a hybrid using the backbone and differential gene expression analysis of ProteoViz package^85^ as a general script workflow and borrowing the normalization script from Proteus package^86^. Briefly, the “proteinGroups.txt” table was read into matrices and filtered for “reverse” hits, “potential contaminant” and proteins “only identified by site”. Data were then normalized using CONSTANd^87^, log2-transformed and differentially analysed using Linear Models for Microarray Data (limma). Significant hits were called proteins with adjusted p-value < 0.005. The final analysis was carried out on combined glucose and galactose samples resulting in 5 replicates for WT and 5 replicates for *Δ*RS. Volcano plots were generated using ggrepel (a ggplot2 extension) as part of the tidyverse. Venn diagram was generated using the VennDiagram package. Scripts for statistical analysis are provided in supplementary data.

## Acknowledgments

GMH acknowledges support from a Faculty of Medicine, Health and Dentistry (FMDH) Research and Innovation Award (312366) from the University of Sheffield, the Royal Society research grant RG140690, the Medical Research Council (MRC) New Investigator research grant MR/R024162/1 and the Biotechnology and Biological Sciences Research Council (BBSRC) grant BB/S005277/1 over the course of this study. SRM was supported by a FMDH PhD studentship funded by the University of Sheffield. HM acknowledges support from a Parkinson’s UK Senior Fellowship (F1301). SKU acknowledges funding from the Francis Crick Institute (grant FC001201), which receives its core funding from Cancer Research UK, the Medical Research Council and the Wellcome Trust.

## Author contributions

GMH designed the overall study. GMH oversaw the biochemical, molecular and cellular biology experiments while the TMT spectrometry and mitochondrial respiratory chain investigations were respectively supervised by SKU and HM. SRM, LMC, YHL, AG, NS and GMH performed the biochemical, molecular and cellular biology assays. SRM, HRF and APS performed the TMT spectrometry. LMC and CH performed the respiratory chain activity assays. SRM undertook is PhD in GMH’s group and is currently a postdoctoral researcher in SKU’s group. GMH and OB supervised the PhD thesis of SRM. GMH wrote the manuscript. SRM, LMC, YHL, HM and SKU edited the manuscript. All authors provided comments and approved the manuscript.

## Conflict of interest

The authors declare no relationships, conditions or circumstances that present a potential conflict of interest.

The authors also confirm that this manuscript has not been published or accepted elsewhere at the time of submission to bioRxiv (19/09/2021).

---

**Supplementary Fig. 1.**
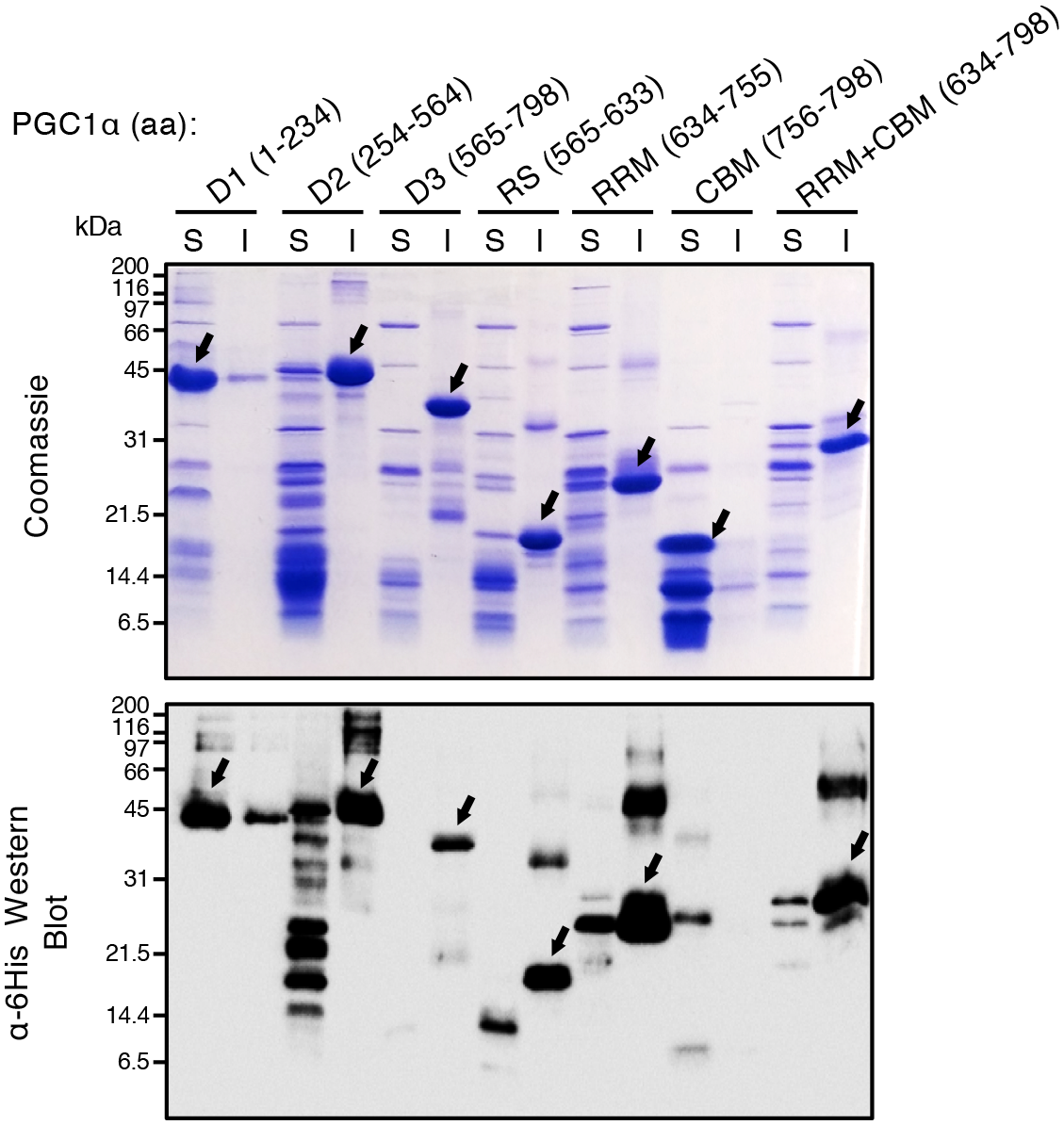
Recombinant expression and purification screening of PGC-1α domains. 6His-tagged protein domains were expressed in *E. coli* BL21 for 3 hours at 37°C prior to small batch purification of soluble (S) and insoluble (I) proteins onto TALON/Cobalt-loaded beads. Eluted proteins were analysed by SDS-PAGE stained with Coomassie blue (top panel) and western blot using α-6His antibodies (bottom panel). Protein domains of PGC-1α are indicated with aa in brackets depicting the amino-acids number positions. 5 µg purified proteins were loaded in each lane. Arrows indicate full length protein domains.

**Supplementary Fig. 2.**
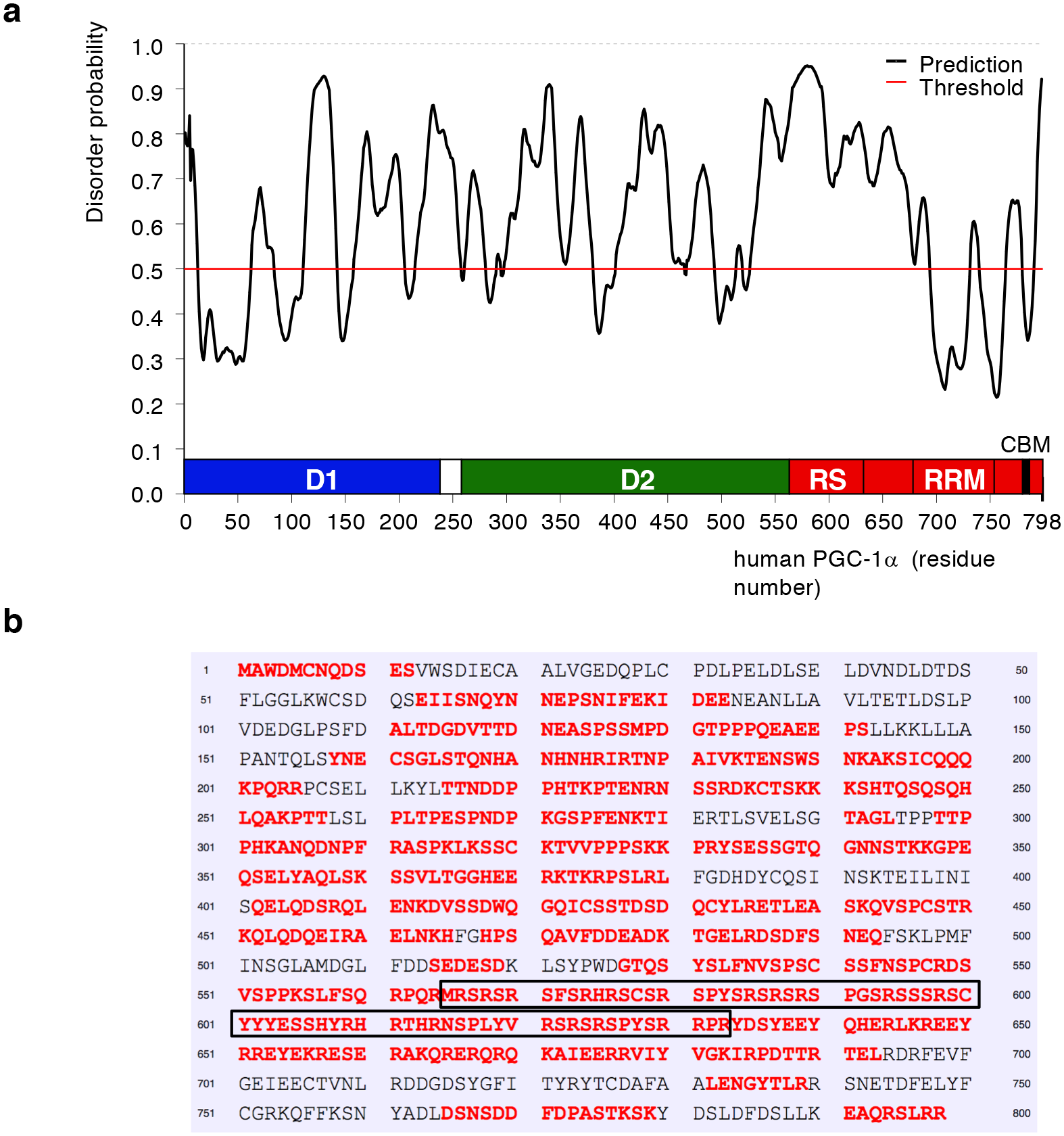
The RS domain of PGC-1α is predicted to be unstructured. **(a)** Probability of disorder is plotted in function of the amino-acid sequence of human PGC-1α using the <u>Pr</u>otein <u>D</u>is<u>O</u>rder prediction <u>S</u>ystem (PrDOS: https://prdos.hgc.jp/cgi-bin/top.cgi) using a prediction false positive rate of 5%). **(b)** Disordered residues are labelled in red. They indicate that a large proportion of the protein is unstructured in agreement with the finding that PGC-1α belongs to the family of intrinsically disordered proteins with a fast turnover mediated by the 20S proteasome (Adamovich *et al*. 2013). The RS domain is highlighted with black boxes (amino acids 565-633).

**Supplementary Fig. 3.**
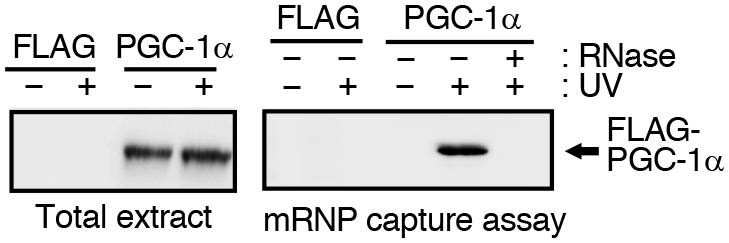
PGC-1α interacts with polyadenylated RNA. Human HEK293T cells were transfected for 48 hours with either FLAG control, FLAG-tagged PGC-1α wildtype or FLAG-tagged PGC-1α ΔRS. Cells were subjected to UV-irradiation (+) or not (-) prior to RNAse treatment of protein extracts (when indicated) and denaturing mRNP capture assays using oligo-dT beads. Eluted proteins were analysed by western blotting probed with α-FLAG antibody.

**Supplementary Fig. 4.**
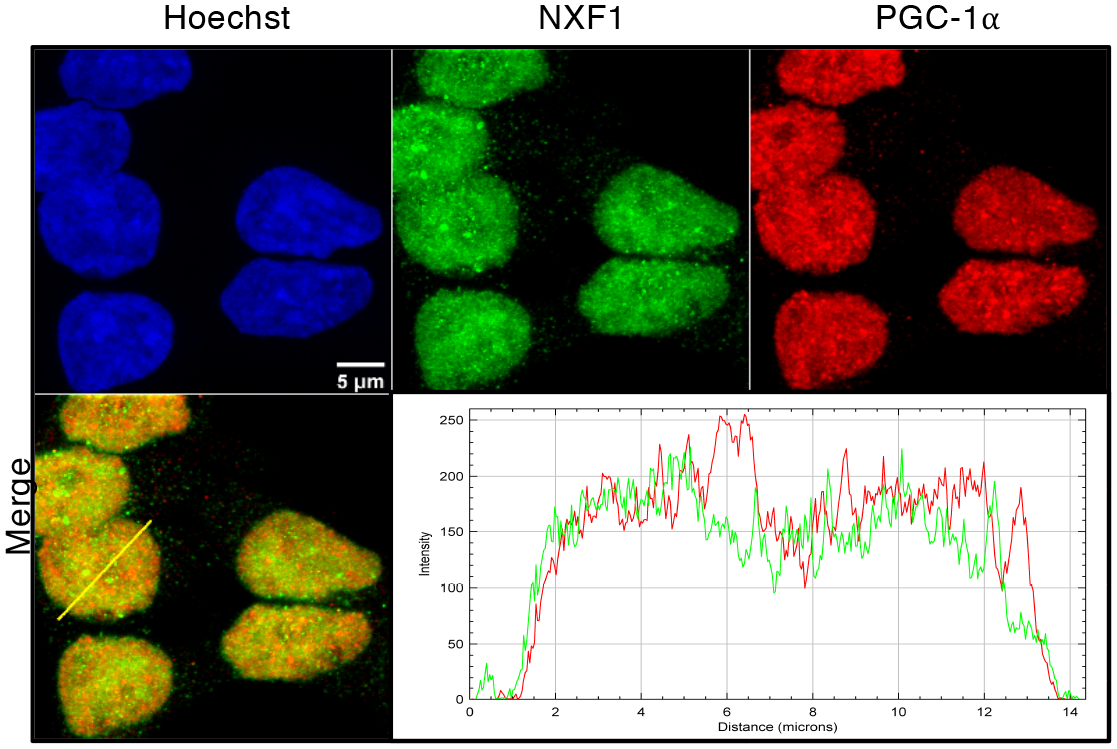
Cellular distribution of endogenous PGC-1α and NXF1 proteins. Confocal immunofluorescence microscopy in HEK293T cells. NXF1 and PGC-1α antibodies were used to labelled endogenous NXF1 and PGC-1α proteins in green and red respectively. Bar scale: 5 µm. Green and red pixel intensities were quantified alongside a nuclear cross-section highlighted with a yellow line.

**Supplementary Fig. 5.**
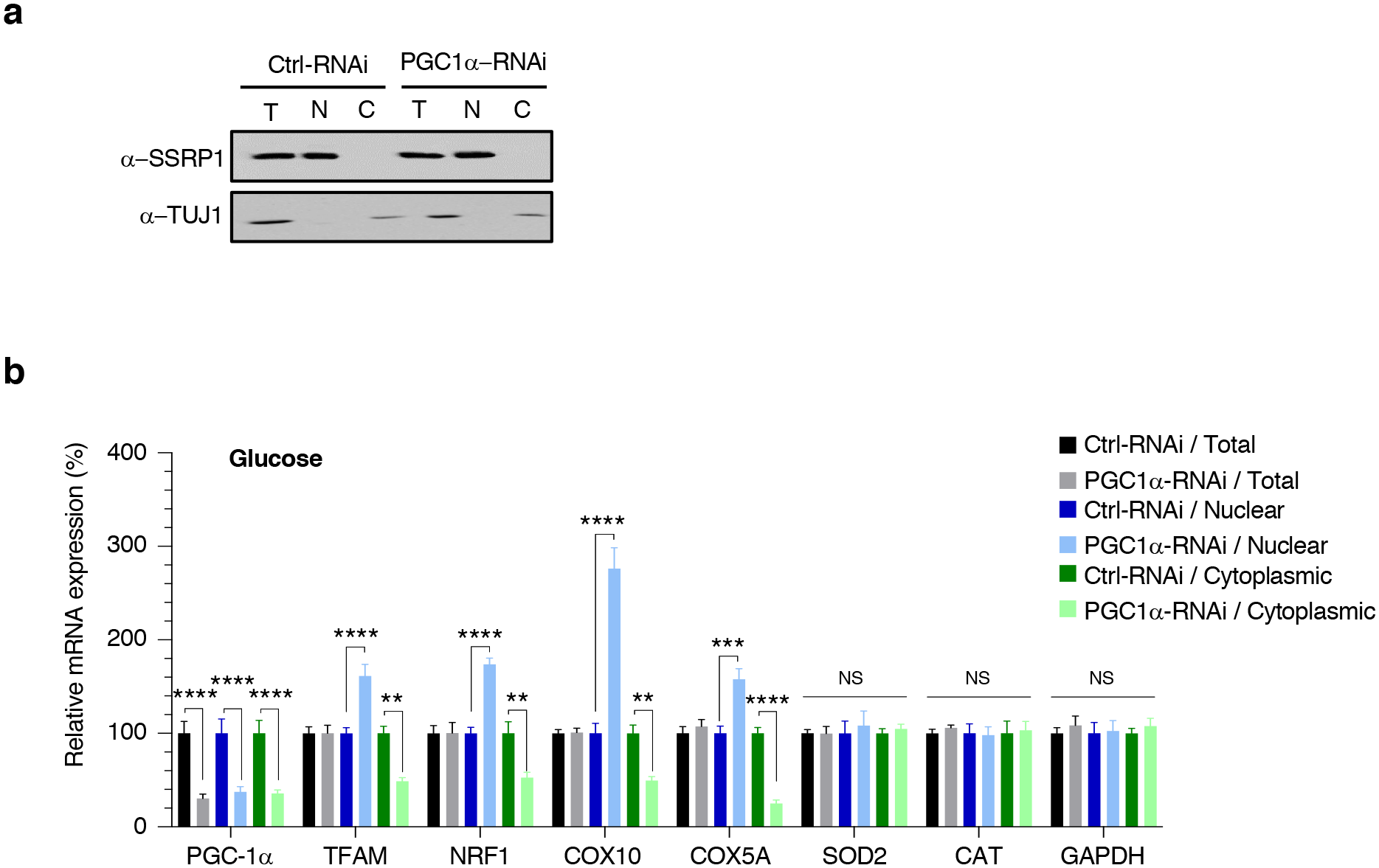
The RS domain of PGC-1α plays drives the nuclear export of mRNAs it transcriptionally co-activates. (**a**) Western blots of Ctrl-RNAi and PGC1α-RNAi cell lines induced for 6 days with doxycycline in glucose-containing medium and subjected to cellular fractionation using hypotonic lysis to yield cytoplasmic fractions. The chromatin remodelling SSRP1 factor is used to check for potential nuclear contamination in cytoplasmic fractions. Depletion of beta-Tubulin III (TUJ1) in nuclear fractions was used to check for quality of the nuclear fractions. **(b)** Total, nuclear and cytoplasmic levels of fractionated RNA were quantified in three biological replicate experiments by qRT-PCR following normalisation to U1 snRNA levels and to 100% in Ctrl-RNAi cell line (Mean ± SEM; two-way ANOVA with Tukey’s correction for multiple comparisons, NS: not significant, **: p<0.01, ***: p<0.001, ****: p<0.0001).

**Supplementary Fig. 6.**
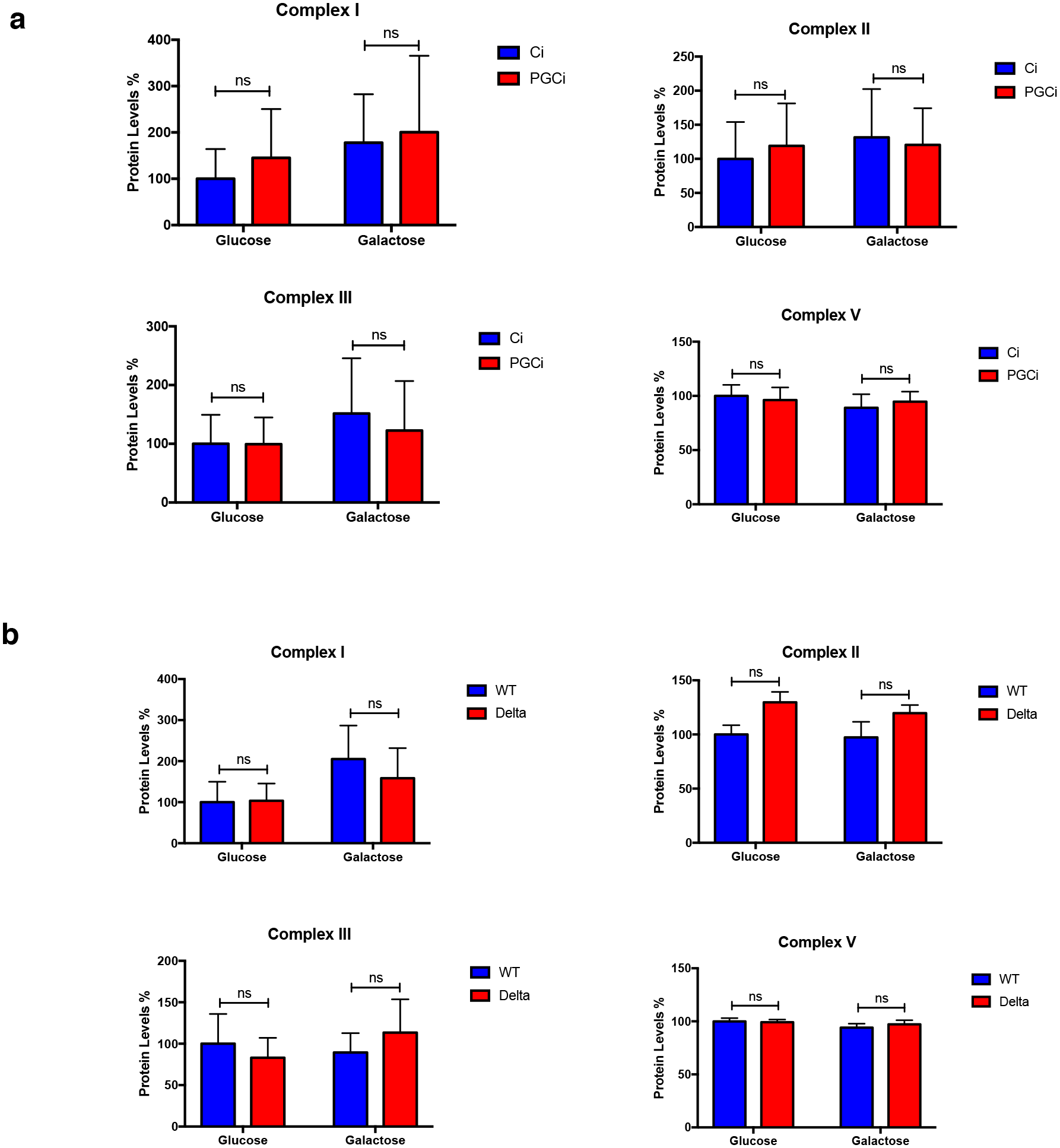
Quantification of OXPHOS western blots in figure 6a-b. **(a)** OXPHOS western blots shown in figure 6a were performed in three biological replicate experiments and changes in protein levels quantified and normalised against the total mitochondrial protein level (the sum of all 5 complexes) (Mean ± SEM; two-way ANOVA with Tukey’s correction for multiple comparisons; NS: not significant). **(b)** OXPHOS western blots shown in figure 6a were performed in three biological replicate experiments and changes in protein levels quantified and normalised against the total mitochondrial protein level (the sum of all 5 complexes) (Mean ± SEM; two-way ANOVA with Tukey’s correction for multiple comparisons; NS: not significant).

**Supplementary Fig. 7.**
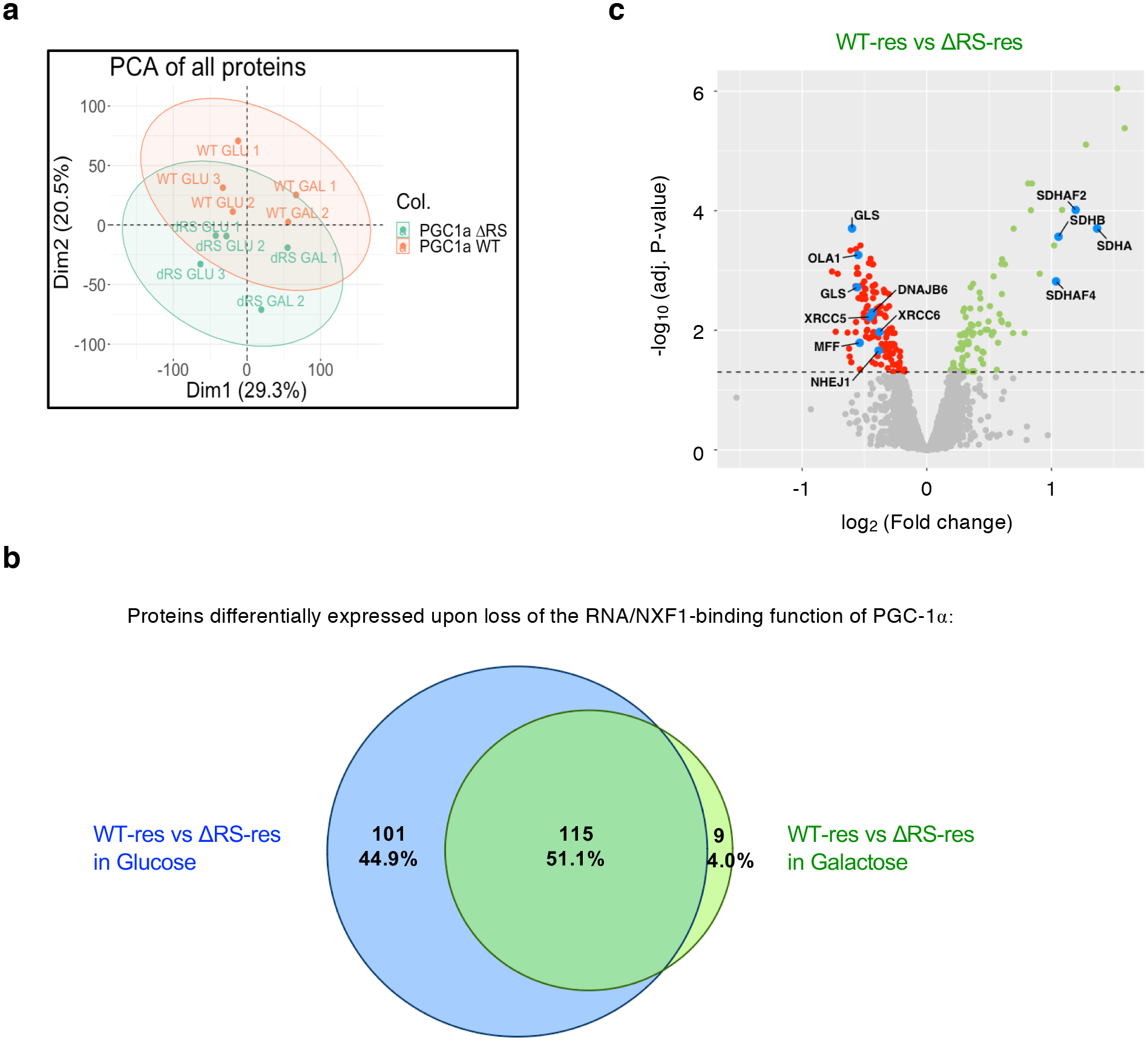
Proteome-wide investigation of the RNA nuclear export of PGC-1α. WT-res and ΔRS-res cell lines were induced with doxycycline for 6 days in media containing glucose and/or galactose for 24 hours prior to proteome identification using Tandem Mass Tag (TMT) spectrometry. **(a)** Principal Component Analysis (PCA) of individual protein samples. **(b)** Venn diagram summarising the identified differential expression of proteins for adjusted p-values <0.05. Numbers (top) indicate dysregulated protein hits while numbers (bottom) highlight the percentage of differentially expressed proteins. **(c)** Volcano plot representing differentially expressed proteins in glucose condition with adjusted p-values <0.05 (n=3 biological replicates).

## SUPPLEMENTARY NOTE 1

### For pcDNA6.2-GW/EmGFP-miR PGC-1α Domain 1

*<u>PGC1a miR D1(133) top:</u>* TGCTGAAAGCTGTCTGTATCCAAGTCGTTTTGGCCACTGACTGACGACTTGGACAGACAGCTTT

*<u>PGC1a miR D1(133) bottom:</u>* CCTGAAAGCTGTCTGTCCAAGTCGTCAGTCAGTGGCCAAAACGACTTGGATACAGACAGCTTTC

### For pcDNA6.2-GW/EmGFP-miR PGC-1α Domain 2

*<u>PGC1a miR D2(997) top:</u>* TGCTGTGTACCAGAAGACTCACTGTAGTTTTGGCCACTGACTGACTACAGTGACTTCT GGTACA

*<u>PGC1a miR D2(997) bottom:</u>* CCTGTGTACCAGAAGTCACTGTAGTCAGTCAGTGGCCAAAACTACAGTGAGTCTTCTGGTACAC

**Quick change oligonucleotides used to confer resistance to chained PGC-1α miRNAs (while not altering the amino-acid sequence of the PGC-1α transgene). Silent mutations are labelled in red:**

<u>PGC miR1 res_fwd:</u>

A CTA GAT GTG AAC GAC TTG GAC ACC GAT TCT TTC CTG GGT GGA CTC AAG TGG TGC

<u>PGC miR1 res_rev:</u>

GCA CCA CTT GAG TCC ACC CAG GAA AGA ATC GGT GTC CAA GTC GTT CAC ATC TAG T

<u>PGC miR2 res_fwd:</u>

A AAG AAG CCC AGG TAC AGT GAA AGC AGC GGC ACC CAA GGC AAT AAC TCC ACC AAG

<u>PGC miR2 res_rev:</u>

CTT GGT GGA GTT ATT GCC TTG GGT GCC GCT GCT TTC ACT GTA CCT GGG CTT CTT T

**Supplementary Note 1 ∣ DNA oligonucleotides used to build the PGC-1α miRNAs and confer resistance to RNAi.**

---

**Supplementary Table 2.** Plasmids used and generated in this study.

| **Plasmid Construct** | **Vector Backbone** | **Tag** | **Cloning Strategy** | **Reference** |
| --- | --- | --- | --- | --- |
| PGC1α 1-798 (FL) | pET24b | N-terminal – GB1 and 6xHis | PCR + NdeI/XhoI | This study |
| PGC1α 1-234 (D1) | pET24b | N-terminal – GB1 and 6xHis | PCR + NdeI/XhoI | This study |
| PGC1α 254-564 (D2) | pET24b | N-terminal – GB1 and 6xHis | PCR + NdeI/XhoI | This study |
| PGC1α 565-798 (D3) | pET24b | N-terminal – GB1 and 6xHis | PCR + NdeI/XhoI | This study |
| PGC1α 565-633 (RS) | pET24b | N-terminal – GB1 and 6xHis | PCR + NdeI/XhoI | This study |
| PGC1α 634-755 (RRM) | pET24b | N-terminal – GB1 and 6xHis | PCR + NdeI/XhoI | This study |
| PGC1α 756-798 (CBM) | pET24b | N-terminal – GB1 and 6xHis | PCR + NdeI/XhoI | This study |
| PGC1α 634-798 (RRM+CBM) | pET24b | N-terminal – GB1 and 6xHis | PCR + NdeI/XhoI | This study |
| PGC1α 1-798 (FL) | pcDNA5 FRT/TO | N-terminal 3xFLAG | PCR + BamHI/NotI | This study |
| Control RNAi | pcDNA5/FRT/TO | miRNA cloned in 3'-UTR EmGFP | BLOCK-iT Kit + PCR + HindIII/NotI | [<sup>51</sup>] |
| PGC1α RNAi (targeting D1 and D2) | pcDNA5/FRT/TO | miRNA cloned in 3'-UTR EmGFP | BLOCK-iT Kit + PCR + HindIII/NotI | This study |
| PGC1α WT res | pcDNA5/FRT/TO | N-terminal 3xFLAG | BamHI/NotI | This study |
| PGC1α ΔRS res | pcDNA5/FRT/TO | N-terminal 3xFLAG | Inverse PCR of WT-res | This study |
| NXF1/TAP | pGEX4T | N-terminal GST | PCR + BamHI/XhoI | [<sup>88</sup>] |
| TAP/NXF1 | pIRES-Neo | C-terminal 13myc | PCR+ BamHI/XhoI | [<sup>40</sup>] |
| p15/NXT1 | pET9a | none | PCR + NdeI/BamHI | [<sup>40</sup>] |
| MAGOH | pET24b | C-terminal 6xHis | PCR + NdeI/XhoI | [<sup>89</sup>] |
| Aly/REF | p3xFLAG | N-terminal 3xFLAG | PCR + EcoRI/XbaI | [<sup>40</sup>] |
| E1B-AP5/hnRNPUL1 | p3xFLAG | N-terminal 3xFLAG | PCR + HindIII/XbaI | [<sup>88</sup>] |
| GFP | p3xFLAG | N-terminal 3xFLAG | PCR + EcoRI/XbaI | [<sup>89</sup>] |

**Supplementary Table 3.**
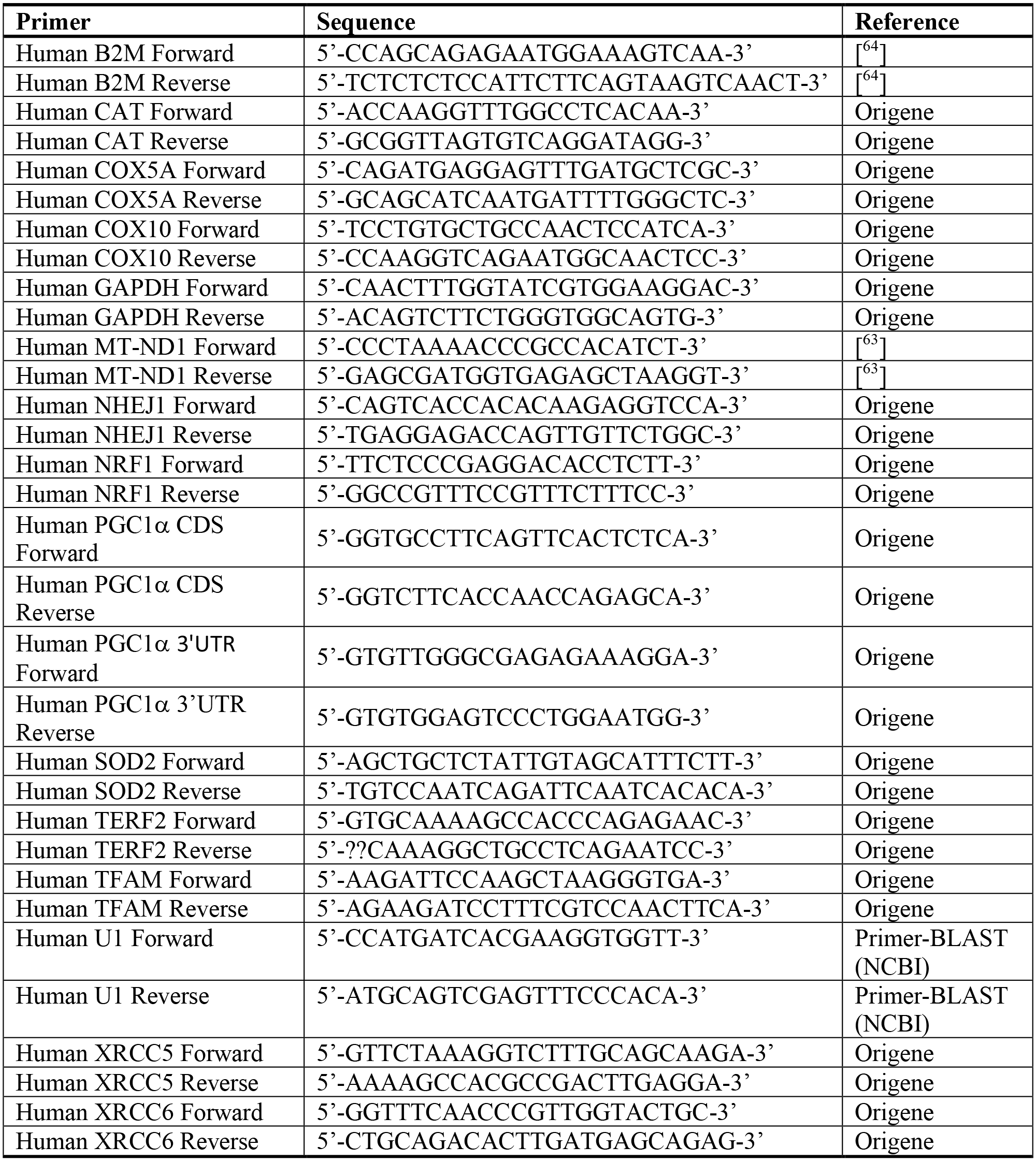
Sequences of qPCR primers used in this study.

